# Competitive dynamics underlie cognitive improvements during sleep

**DOI:** 10.1101/2021.03.31.437952

**Authors:** Pin-Chun Chen, Hamid Niknazar, William A. Alaynick, Lauren N. Whitehurst, Sara C. Mednick

**Affiliations:** Department of Cognitive Sciences, University of California, Irvine; Department of Statistics, University of California, Irvine; ScholarNexus, LLC; Department of Psychology, University of Kentucky

**Keywords:** working memory, long-term memory, sleep, vagal activity, spindle activity

## Abstract

We provide evidence that human sleep is a competitive arena where cognitive domains vie for limited resources. Using pharmacology and effective connectivity analysis, we demonstrate that long-term memory and working memory are served by distinct offline neural mechanisms that are mutually antagonistic. Specifically, we administered zolpidem to increase central sigma activity and demonstrated targeted suppression of autonomic vagal activity. With effective connectivity, we determined the central activity has greater causal influence over autonomic activity, and the magnitude of this influence during sleep produced a behavioral trade-off between offline long-term and working memory processing. These findings show the first evidence of a sleep switch mechanism that toggles between central sigma-dependent long-term memory and autonomic vagal-dependent working memory processing.

**Significant Statement:** Sleep facilitates both long-term episodic memory consolidation and short-term working memory functioning. However, the mechanism by which the sleeping brain performs both complex feats, and which sleep features are associated with these processes remain unclear. Using a pharmacological approach, we demonstrate that long-term and working memory are served by distinct offline neural mechanisms, and that these mechanisms are mutually antagonistic. We propose a Sleep Switch model in which the brain toggles between the two memory processes via a complex interaction at the synaptic, systems, and mechanistic level, with implications for research on cognitive disturbances observed in neurodegenerative disorders such as Alzheimer’s and Parkinson’s disease, both of which involve the decline of sleep.

## Main Text

Working memory (WM) and long-term memory (LTM) serve separate functions and the idea that they are supported by separate systems has become a core assumption of modern cognitive psychology [1]. WM is a control process for planning and carrying out behavior that is information-independent, whereas LTM is an information-dependent vast store of knowledge and record of prior events. Both WM and LTM rely on offline periods that include sleep to facilitate performance improvement. According to the framework of systems consolidation, long-term memories are initially bound by a fast-learning system in the hippocampus (i.e. encoding), followed by stabilization of these memory traces in cortical stores (i.e., consolidation). Non-rapid-eye-movement (NREM) sleep may facilitate consolidation by increasing communication between cortico-thalamo-hippocampal circuits via nested oscillations of slow oscillations (<1Hz, SO), spindles (sigma power; 12-15Hz), and sharp wave ripples (SPW-R), respectively [2–4]. SOs reflect fluctuations of the membrane potential and orchestrate transitions from neuronal silence (hyperpolarized downstates) to neuronal excitation (depolarized upstates). Spindles, nested in SO upstates, gate dendritic Ca2+ influx and promote synaptic plasticity. Hippocampal SW-Rs nested in spindles are closely linked to the reactivation of cell assemblies engaged during encoding. Prior studies suggested that spindles may initiate hippocampal-cortical dialogue by grouping SW-Rs, which facilitates information transfer between neocortical and hippocampal cell assemblies. In humans, pharmacological interventions that boost spindle activity enhance sleep-dependent hippocampal LTM, measured by the paired-associates task [5–7].

Classic models of WM propose two governing mechanisms: 1) an active maintenance of information online through the elevated firing of prefrontal neurons, and 2) a supervisory executive control process that is supported by a prefrontal-subcortical inhibitory network [8, 9]. Due to innervations to the heart via sympathetic stellate ganglia and parasympathetic vagal nerve efferents, cardiac autonomic activity is thought to reflect functioning of prefrontal inhibitory processing [10]. Accordingly, vagally-mediated, high frequency heart rate variability (0.15-0.40, HF HRV) during wake correlates with executive function tasks, such as WM, which rely on PFC activity [11]. Improvement in WM, however, only occurs when the interval between training sessions contains a period of sleep, measured by N-back, complex-span task, and digit span [12–17]. Although the exact mechanisms of WM improvement during sleep are still not entirely understood, prior studies point to SWS as an optimal state for synaptic plasticity and cortical reorganization. During SWS, vagal activity is also at its highest compared to all other states of consciousness [18]. Building on this foundation, a recent study identified vagal HF HRV during SWS as a strong predictor of WM improvement, measured by the operation-span task [19].

Together, theoretical models and empirical data suggest that NREM sleep may facilitate improvement in WM via strengthening of prefrontal-autonomic inhibitory networks, measured by HF HRV, while facilitating the formation of LTM via thalamic spindles driving the hippocampal-cortical dialogue, measured by sigma power. The question is how the sleeping brain performs both of these complex feats and which sleep features are associated with these processes? Prior animal studies suggest a potentially antagonistic interplay between the cortico-thalamo-hippocampal networks and the prefrontal-autonomic inhibitory networks [20, 21]. However, this possibility and its functional significance has not been studied in humans.

In the present study, we enacted a pharmacological strategy to investigate the bi-directional interplay between central (reflected in sigma activity) and autonomic (reflected in vagal HRV) activities during overnight sleep and its impact on LTM and WM, measured by the word-paired associative task and the operation-span task. Specifically, we tested our model that central sigma activity would suppress autonomic vagal activity using effective connectivity [22], defined as the influence that one neural system exerts over another, which can be estimated using Granger causality (Fig. 3a). We identified a novel antagonistic relationship between sigma and vagal activity during sleep, with the degree of mutual antagonism between sigma and vagal activity predicting a heretofore unreported behavioral trade-off between LTM and WM. These results suggest that NREM sleep confers benefits to WM and LTM by switching between separate offline mechanisms, i.e., the prefrontal-autonomic inhibitory processing and the hippocampal-cortical dialogue. Furthermore, this sleep switch can be biased towards LTM consolidation by increasing sigma activity, in this case pharmacologically, and presumably by other methods as well. These results illuminate the dynamics interplay underlying LTM and WM processes during sleep.

## Results

### Experiment 1

Based on previous findings, we predicted that central sigma power would have an inhibitory effect on cardiac vagal tone. To this end, we administered zolpidem in a double-blind, placebo-controlled, randomized cross-over design, in which each participant experienced two nights per drug condition (zolpidem or placebo; a total of 4 nights; *N* = 34; *M*_age_ = 20.88 ± 1.88 years, 17 Females), with EEG and ECG monitored (Fig. 1 shaded area). The order of drug conditions was counterbalanced with at least a one-week interval between the experimental visits to allow for drug clearance. We performed power spectral analysis to quantify normalized sigma activity and analyzed HRV profiles. Our intervention was successful, whereby zolpidem increased time spent in SWS while decreasing WASO (*Supplemental Table. S2*), and enhanced sigma activity during stage 2 sleep (central channels: t = 2.112, p = .0349; parietal channels: t = 2.214, p = .0270, corrected by Tukey’s multiple comparisons; *Supplemental Table. S7*), consistent with prior literature.

**Fig. 1.**
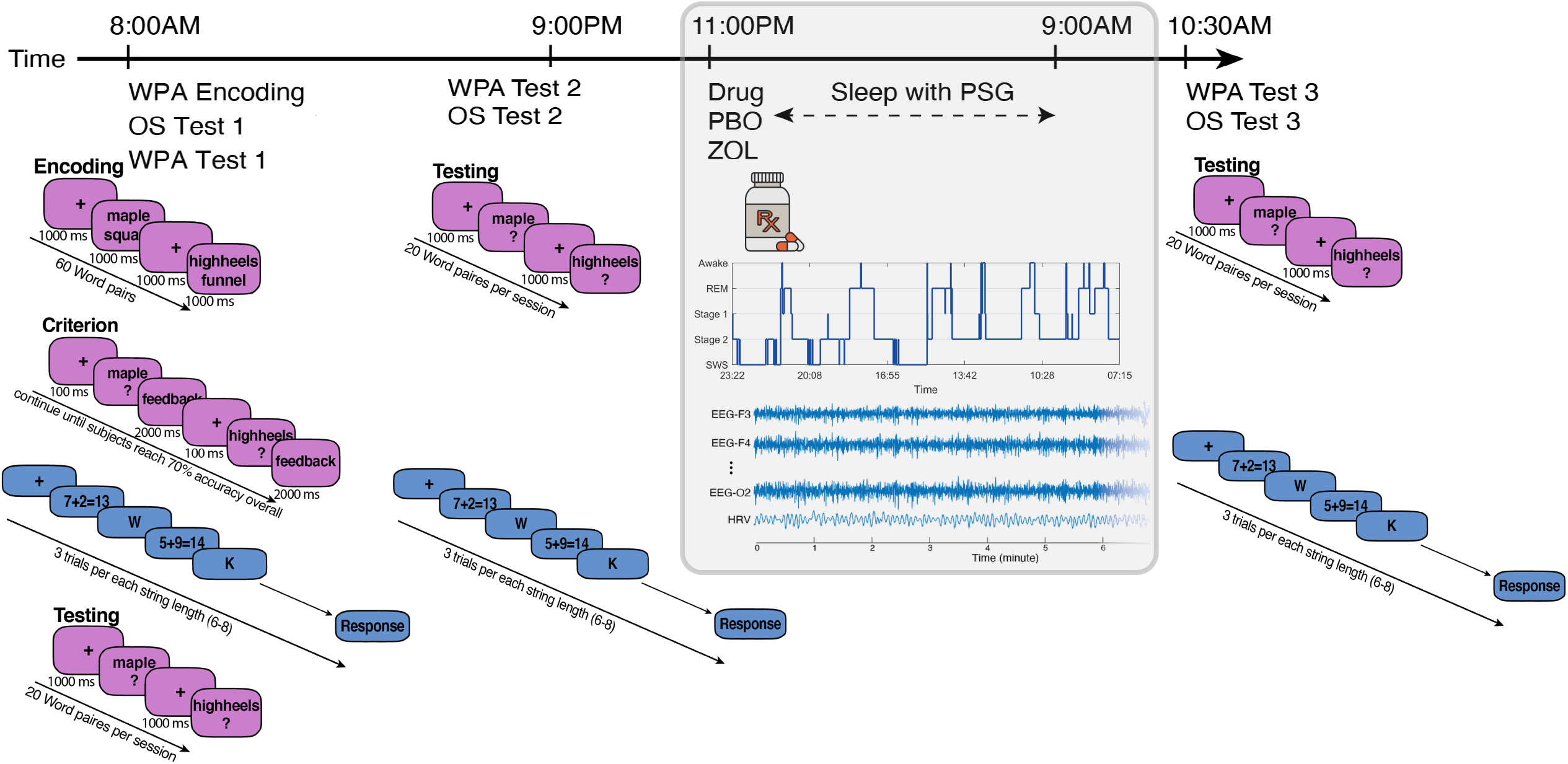
Experimental design and behavioral tasks. **Experiment 1**: Participants reported to the lab at 9:00PM and were hooked up to polysomnography (PSG), including electroencephalography (EEG), electrocardiogram (ECG), electromyogram (EMG), and electrooculogram (EOG). Before sleep, we recorded 5-min resting HRV while subjects lay awake in a still, supine position. At 11:00PM, directly before lights-out, subjects ingested either 10mg of zolpidem or placebo. Sleep was monitored online by a trained sleep technician. Participants were woken up at 9:00AM the next morning and permitted to leave the lab. Each participant experienced two visits per drug condition (a total of four visits). **Experiment 2**: At 8:00AM, participants began encoding for the episodic memory word-paired-associates (WPA) task, followed by the working memory operation-span task (OS) task and immediate recall for the WPA (Test 1). Participants left the lab after cognitive testing. Participants were asked not to nap, exercise, or consume caffeine or alcohol, and were monitored with actigraphy during the break. Participants returned to the laboratory at 9:00 PM to complete the delayed recall over wake for WPA and OS (Test 2). Participants were then hooked up to polysomnography (PSG), including electroencephalography (EEG), electrocardiogram (ECG), electromyogram (EMG), and electrooculogram (EOG). Before sleep, we recorded 5-min resting HRV while subjects lay awake in a still, supine position. At 11:00PM, directly before lights-out, subjects ingested either 10mg of zolpidem or placebo. Sleep was monitored online by a trained sleep technician. Participants were woken up at 9:00AM the next morning and provided a standardized breakfast. At 10:30 AM, participants completed the delayed recall over sleep for WPA and OS (Test 3). For both tasks, to assess the change in performance, we measured two difference scores: overnight change (Test 3 – Test 2); 24-hr change (Test 3 – Test 1). Each participant experienced one visit per drug condition (a total of two visits). See *Supplemental Fig. S1 and Table S12* for summary statistics.

As we hypothesized, zolpidem not only increased sigma activity, but also selectively suppressed vagal tone during sleep, measured by root mean square of the successive differences (RMSSD) (Fig. 2a) and high-frequency HRV (HF; Fig. 2b), but had no impact on low-frequency HRV (0.04-0.15, LF; Fig. 2c). Other HRV indices were reported in *Supplemental Fig. S2 and Table S4*.

**Fig. 2.**
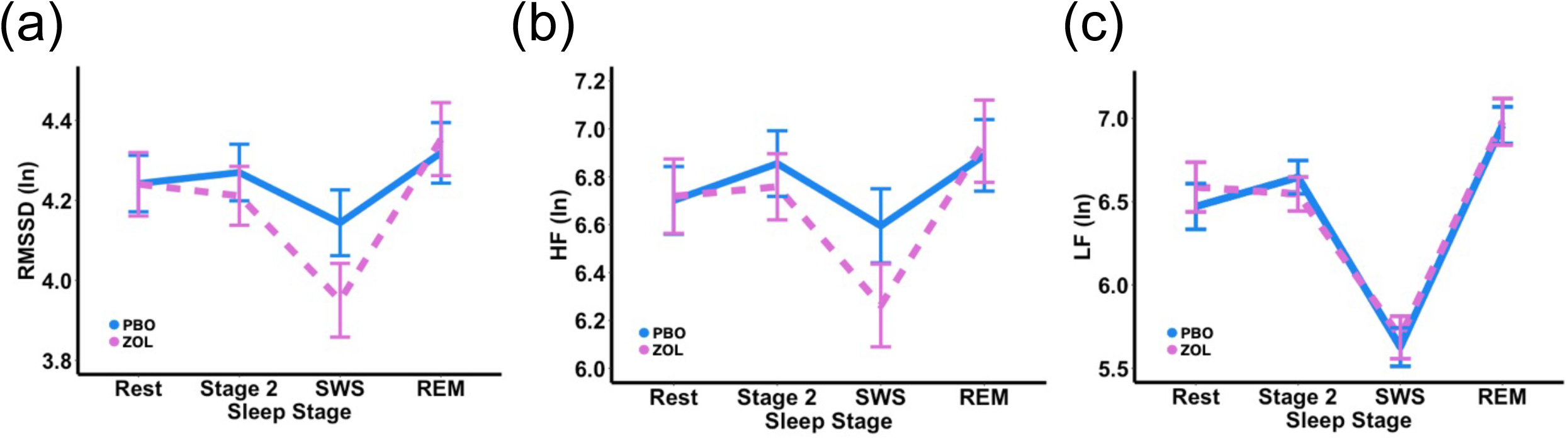
Zolpidem decreased vagally-mediated HRV, but not LF, during SWS. **(a) RMSSD:** We report a significant main effect of sleep stage (F(3, 366) = 21.257, p < .0001), with a decreased HRV during SWS compared to Rest, Stage 2, and REM (all ps < .0001). We also found a significant interaction (F(3, 366) = 3.8630, p = .0096) between sleep stage and drug condition, with decreased vagal activity during SWS (p = .0006) in zolpidem compared with placebo, but not during Stage 2 (p = .3549), REM (p = .3804), or Rest (p = .6152). The likelihood ratio test was significant (LR = 13.8544; p = .0078), suggesting that zolpidem significantly modulated the time-domain measure of HRV. **(b) High-frequency (HF) HRV:** We report a significant main effect of sleep stage (F(3, 366) = 16.9891, p < .0001), with a decreased HRV during SWS compared to Rest (p = .0006), Stage 2 (p < .0001), and REM (p < .0001). Similarly, we also report a significant interaction (F(3, 366) = 3.1899, p = .0238) between sleep stage and drug condition, with decreased vagal activity during SWS (p = .0020) in zolpidem compared with placebo, but not during Stage 2 (p = .4194), REM (p = .4365), or Rest (p = .6070). The likelihood ratio test was significant (LR = 11.3671; p = .0227), suggesting that zolpidem significantly modulated the frequency-domain measure of HRV. **(c) Low-frequency (LF) HRV:** We report a significant main effect of sleep stage (F(3, 366) = 93.0330, p < .0001), with a decreased LF power during SWS compared to Rest, Stage 2, and REM (all ps < .0001), and an increased LF power during REM compared to Rest and Stage 2 (all ps < .0001). No significant main effect of drug condition (p = .6337), nor interaction between sleep stage and drug condition (p = .5681) were found. The likelihood ratio test was not significant (LR = 2.2889; p = .6828), suggesting that zolpidem did not significantly modulate low frequency HRV.

We then tested our hypothesis that central sigma power would exert greater causal influence over vagal autonomic activity than the influence of vagal over sigma activity, and such difference would be increased by zolpidem. To test this prediction, we used effective connectivity estimation (Fig. 3a). In particular, we tested the hypotheses that central sigma naturally exercises greater causal influence on autonomic vagal activity than vice versa in the placebo condition, and that increasing sigma with zolpidem would increase causal information flow from sigma to vagal activity, while decreasing the causal information flow from vagal to sigma activity in the zolpidem condition. For each subject, we calculated two measures: HFInflow and HFOutflow, respectively (see Methods). We confirmed our hypothesis that central sigma power exerted greater flow on vagal activity than vice versa in the placebo condition (p < .0001; Fig 3b). We also confirmed that such difference was increased by zolpidem (p = .0369; Fig. 3b). Next, we calculated a composite score, the effective connectivity ratio: HFInflow over HFOutflow, where higher numbers represented greater central sigma control over autonomic vagal activity. We observed a higher effective connectivity ratio during the zolpidem night (p = .0059). Taken together, results from Experiment 1 were consistent with our hypotheses that central sigma activity naturally exerts dominance over autonomic activity during NREM sleep, and that increasing sigma activity via zolpidem inhibits vagal activity and enhances central sigma control over autonomic vagal activity.

**Fig. 3.**
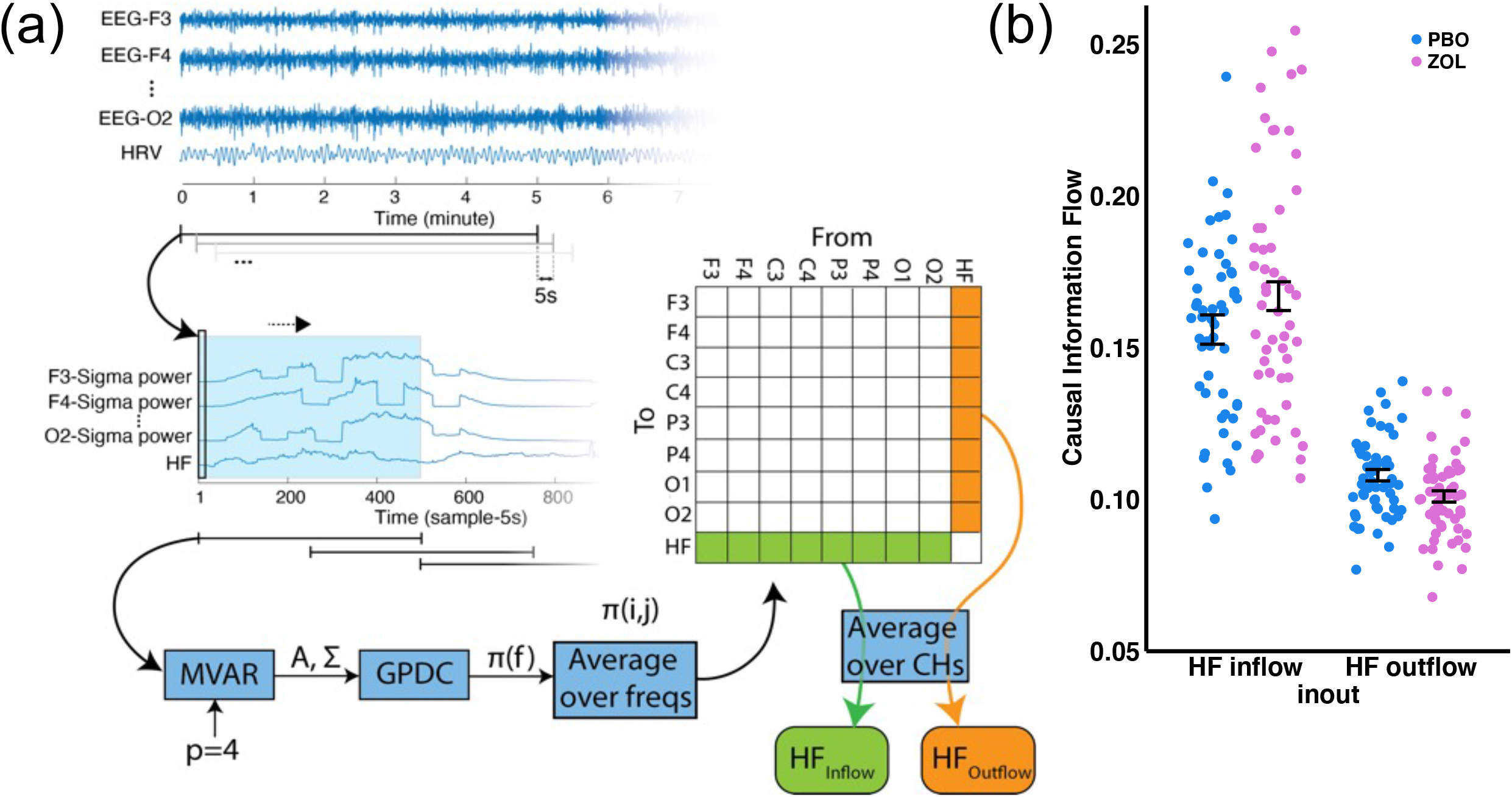
Effective Connectivity Modulated by Drug Condition. **(a) Effective Connectivity Estimation Procedure** (see Methods for details): Prior research using functional connectivity analysis has measured temporal similarity or correlations between different EEG channels [53]. Although functional connectivity can reveal important information about communication, it is limited to correlational measures, and cannot identify directional causal communication. In contrast, Effective connectivity is defined as the influence that one neural system exerts over another either directly or indirectly [22], which can be estimated using Granger causality [54]. According to Granger causality, a causal relation is detected if past values of a source signal help predict a second signal (sink signal) beyond the information contained in its past alone. Granger causality and causal information flow can be quantified using a multivariate vector autoregressive model (MVAR) and then examining the coefficients of the fitted model. Partial directed coherence (PDC) quantifies direct causal information outflow from each signal to all other signals, emphasizing the sinks, rather than the sources [55]. The current study adopted the generalized form of PDC (GPDC) to quantify causal information flow [56] with respect to both the source and the sink regions. Model order (p) of the MVAR model was the only parameter and was selected based on the Akaike information criterion (AIC). **(b) Experiment 1 Effective Connectivity:** We report a main effect of inflow vs outflow (F(1, 185) = 273.317, p <.0001), with a greater HFinflow than HFoutflow in both drug conditions; an interaction between drug condition and inflow vs outflow (F(1, 185) = 5.744, p = .0175), with a greater HFinlfow during zolpidem compared to placebo (p = .0369). No main effect of drug condition was found (F(1, 185) = 0.512, p = .4751). The likelihood ratio test was significant (LR = 6.0745; p = .0480), suggesting that zolpidem significantly modulated the causal information flow between sigma and HF activity. Effective connectivity ratios (HFInflow/ HFOutflow) increased significantly during the zolpidem night (F(1, 79) = 8.0607, p = .0059).

### Experiment 2

In an independent sample of participants (*N* = 38; *M*_age_ = 20.85 ± 2.97 years; 19 Females), we added a behavioral experiment (Experiment 2; Fig. 1) to the original design of Experiment 1 to test whether we could replicate the physiological results of Experiment 1 and determine their functional importance for sleep-dependent cognition. Again, we exploited zolpidem to modulate the interaction between central sigma and autonomic vagal activity and examined its impacts on the improvements of LTM and WM (Fig. 1). The order of drug conditions was counterbalanced with at least a one-week interval between the two experimental visits to allow for drug clearance. The goal of experiment 1 was to thoroughly describe the physiological phenomenon across the whole night, whereas the goal for experiment 2 was to examine the functional impact of the pharmacological intervention on performance. For this reason, in experiment 2 we divided the night into quartiles and focused our analyses on quartile two and three combined to maximize zolpidem’s effect, due to the pharmacodynamics of zolpidem, which has a half-life of (1.5–4.5 h), and onset (mean Tmax 1.6h [23]). We hypothesized that sigma-guided vagal suppression effects would result in parallel behavioral effects with greater long-term memory and reduce improvement in working memory. We further hypothesized that the magnitude and the direction of causal information flow between central and autonomic systems would be correlated with the trade-off between LTM and WM.

The physiological results across one night of sleep in Experiment 2 were consistent with those from two nights of sleep in Experiment 1 (see *Supplemental Table S3 for sleep architecture*; *Supplemental Table S8 for power spectrum; Supplemental Fig. S3, S4, S5, S6, Table S5, S6 for HRV; Supplemental Fig. S7 for effective connectivity)*. We confirmed that zolpidem increased sigma activity during sleep while suppressing vagal tone, measured by RMSSD and HF, but had no impact on LF. Similarly, we replicated the effective connectivity results (Fig. 3c), in which zolpidem increased effective connectivity ratio (p = .0265), indicating greater causal influence of central sigma activity on autonomic vagal activity.

We further assessed the functional roles of each physiological measure (EEG sigma activity, cardiac vagal activity, and effective connectivity ratio) on LTM and WM changes across sleep. We hypothesized that increasing sigma activity would benefit LTM retention in a word-pair-associates (WPA) task, whereas decreasing vagal activity would hinder WM improvement on a working memory operation span (OS) task. To this end, we examined overnight and 24-hr change scores in each task between the two drug conditions. For the word-pair task, our analysis showed that zolpidem significantly increased 24-hr LTM retention (Fig. 4a right panel) and overnight retention (Fig. 4a left panel). For the working memory operation span task, our analysis demonstrated that zolpidem decreased overnight improvement (Fig. 4b left panel) and 24-hr improvement (Fig. 4b right panel), compared to placebo. In summary, we confirmed our behavioral hypothesis that sigma-guided vagal suppression would increase LTM (Fig. 4a) and decrease WM improvement (Fig. 4b).

**Fig. 4.**
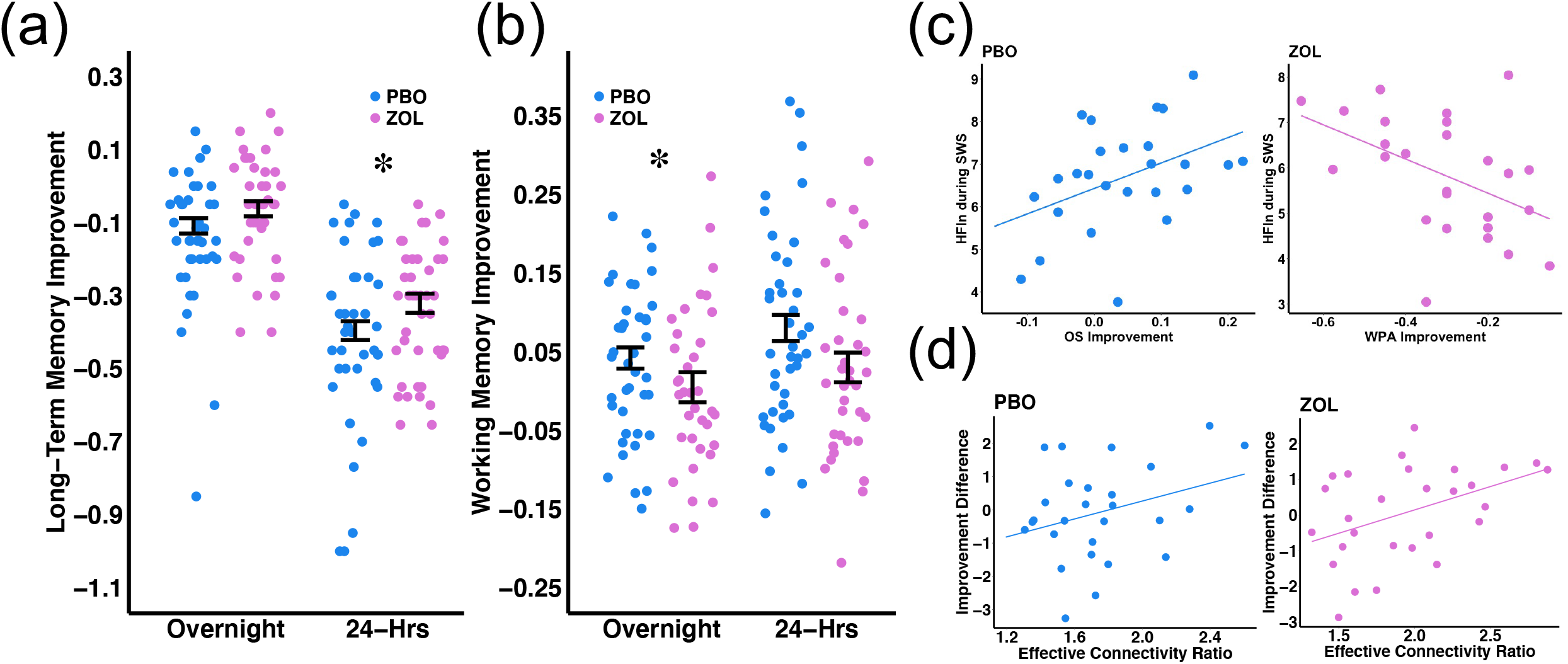
Zolpidem increases LTM but decreases WM improvement. **(a) Long-term memory (WPA task) improvement by drug conditions and time.** (Y axis: WPA Overnight [Test3-Test2] and 24-hr [Test3-Test1] improvement; asterisks indicate significant differences in behavioral changes between two drug conditions; *p<0.05) ZOL yielded greater but not significant overnight retention of WPA than the PBO condition (estimate= −0.1156, CI= (−0.2408, −0.0095), t= −1.8104, p= 0.0810), accounting for visit, as well as greater 24-hr retention of WPA than PBO visits (estimate= −0.1810, CI= (−0.3519, −0.0096), t= −2.0704, p= 0.0474), accounting for visit. **(b) Working memory (OS task) improvement by drug conditions and time.** (Y axis: OS Overnight [Test3-Test2] and 24-hr [Test3-Test1] improvement; asterisks indicate significant differences in behavioral changes between two drug conditions; *p<0.05) PBO showed significantly greater overnight improvement of OS than ZOL visits (estimate= 0.1242, CI= (0.0201, 0.2284), t= 2.3377, p= 0.0260), accounting for Test 2 performance and visit, as well as greater but not significant 24-hr improvement of OS than ZOL visits (estimate=0.1000, CI= (−0.0184, 0.2185), t= 1.6546, p= 0.1081), accounting for Test 1 performance and visit. **(c) Functional role of vagal activity on memory.** (Y axis: HFln during SWS, X axis: OS overnight and WPA 24-hr improvement) Vagal activity during SWS positively predicted working memory (OS task) improvement (r = .422; p = .032) but negatively predicted long-term memory (WPA task) improvement (r = −.460; p = .018). The difference between these two correlations was significant (Z = 3.67; p = .0001). **(d) Functional role of effective connectivity ratio on memory trade-off.** (Y axis: normalized WPA improvement - normalized OS improvement score, X axis: effective connectivity ratio = HFInflow/ HFOutflow) Effective connectivity ratio (a higher ratio indicates a greater causal effect from sigma to vagal) during sleep positively predicted memory trade-off (a greater difference indicates a greater improvement in the WPA task than the OS task) during the zolpidem night (r = .429; p = .020), but not the placebo night (r = .251; p = .190). The difference between these two correlations not significant (Z = 0.78; p = 0.2177).

Next, we tested the correlations between each physiological measure (EEG sigma activity, cardiac vagal activity, and effective connectivity ratio) and memory changes across sleep using Pearson’s correlation coefficients. We found a functional dissociation in vagal activity and behavior, where vagal activity during SWS was negatively correlated with LTM in the zolpidem condition (24-hr retention and HFln: r = −.460; p = .018; Fig. 4c right panel), and positively correlated with WM improvement (overnight retention and HFln: r = .422; p = .032; Fig. 4c left panel) in the placebo condition. We compared correlations between HFln and LTM versus HFln and WM, and the difference was significant (Z = 3.67; p = 0.0001). This result is in line with our expectation that vagal activity during sleep differentially supports LTM and WM. Correlational statistics between vagally-mediated HRV parameters and behavioral improvements are shown in *Supplemental Table S9*. No significant correlations were found between EEG sigma activity and WM improvement (zolpidem: all ps > .5687; placebo: all ps > .1943) or between EEG sigma activity and LTM retention (zolpidem: all ps > .15516; placebo: all ps > .1383; see *Supplemental Fig. S8 for spindle density*). Taken together, vagal activity was positively associated with WM improvement, but inversely related to LTM.

We, then, asked whether central and autonomic antagonism impacted the trade-off between LTM and WM improvement by correlating the effective connectivity ratio with the normalized LTM-WM difference score, where higher numbers represent greater LTM than WM improvement. We found a positive correlation between the effective connectivity ratio and normalized LTM-WM difference score in the zolpidem (r = .429; p = .020; Fig. 4d right panel) and non-significant positive correlation in the placebo condition (r = .251; p = .190; Fig. 4d left panel). These results suggested that the more central activity exerted influence on autonomic vagal activity, the more sleep was biased towards sigma-dependent LTM consolidation (and away from vagal-dependent WM processing). We further compared correlations between LTM-WM difference score and the effective connectivity ratio in the placebo versus zolpidem condition. The difference was not significant (Z = 0.78; p = 0.2177), suggesting that zolpidem amplified the natural vagal suppression by sigma and thus increased the magnitude of the correlations.

Given the critical role for system consolidation of nested oscillations between sigma and SOs, the current findings led us to the prediction that greater sigma-SO coupling would evince increased LTM via suppressed WM. We tested this prediction by computing sigma power during the up-state of SOs and correlating this magnitude with the normalized LTM-WM improvement difference score *(see Supplemental Table S10 for SO counts and Sigma/SOs Summary Statistics; Supplemental Table S11 for correlations)*. We found that zolpidem decreased the number of SOs, a finding consistent with prior literature that zolpidem shifts brain activity to faster frequencies. This decrease in SOs by zolpidem led us to examine coupling in the placebo condition, in which we found a significant positive correlation between sigma power during SOs up-state and difference in LTM-WM improvement (Fig. 5), consistent with the notion that competitive dynamics underlie the fundamental mechanisms of cognitive improvements during sleep.

**Fig. 5.**
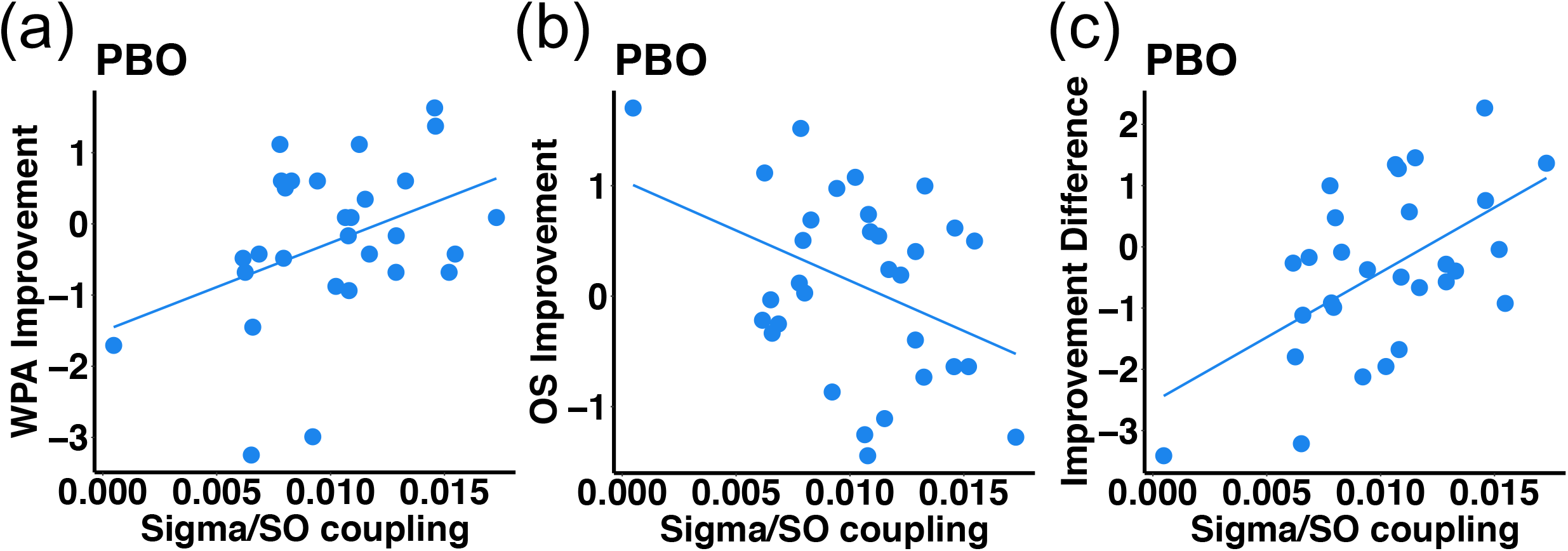
Functional roles of sigma power coupled with SO up-state on LTM and WM. **(a) Long-term memory (WPA task) improvement positively correlated with sigma power coupled with SO up-state.** (Y axis: normalized score of WPA 24-hr improvement; X axis: normalized sigma power coupled during the up-state of SOs; r = .400; p = .034) **(b) Working memory (OS task) improvement negatively correlated with sigma power coupled with SO up-state.** (Y axis: normalized score of OS overnight improvement; X axis: normalized sigma power coupled during the up-state of SOs; r = −.380; p = .033) **(c) Improvement difference positively correlated with sigma power coupled with SO up-state.** (Y axis: normalized WPA improvement - normalized OS improvement score; X axis: normalized sigma power coupled during the up-state of SOs; r = .560; p = .002)

## Discussion

The current work identified two neural mechanisms during NREM sleep that support the distinct enhancements in long-term and working memory. In experiment 1, we exploited the hypnotic zolpidem to enhance sigma activity during NREM sleep and report the novel finding that increasing sigma activity resulted in targeted vagal suppression during NREM. Next, we used the effective connectivity estimation technique to test the causal hypothesis that central sigma activity actively suppressed vagal autonomic activity. Consistent with our hypothesis, results showed that central sigma exerted greater causal control over autonomic vagal activity and that pharmacologically increasing sigma activity boosted causal information flow from central to autonomic channels and decreased flow from autonomic to central channels. In a separate set of subjects, we replicated the pharmacological intervention and tested the functional significance of the sigma-vagal mutual antagonism during NREM sleep by testing LTM and WM before and after a night of sleep. The physiological and effective connectivity results replicated those of experiment 1. Moreover, the sigma-guided vagal suppression was associated with enhanced LTM retention at the cost of reduced WM improvement. Additionally, the magnitude of vagal suppression, as well as the degree of sigma-SO coupling, predicted a not previously reported trade-off between LTM and WM processing. These findings suggest evidence for a sleep switch that toggles between separate and non-overlapping NREM mechanisms that support LTM and WM processing. Furthermore, this switch can be biased towards greater LTM consolidation by boosting sigma activity.

Sigma activity is proposed to facilitate plasticity by producing long-term changes in responsiveness in cortical neurons [24] and increasing dendritic Ca^2+^ influxes [25], particularly enhanced when coupled to down-to-up transitions of the sleep slow oscillation. Recently, Dickey and colleagues demonstrated sigma activity may promote spike-timing-dependent plasticity (STDP), which facilitates long-term potentiation (LTP), the cellular mechanism thought to underlie learning and memory [26]. Thus, sigma activity may promote LTM via cellular synaptic plasticity. Furthermore, at the systems level, sigma nested within SOs may also support the replay of memory traces during consolidation [27], and causally increasing sigma activity boosts hippocampal-dependent memory consolidation [5, 28, 29]. The current findings demonstrate that sigma activity, especially when coupled with SOs also suppresses subcortical vagal activity with significant functional outcomes, specifically a reduction in WM.

Vagal influence on cognitive function is a core principle of the Neurovisceral Integration Model [10], which posits that ANS activity is a peripheral index of the integrity of prefrontal-autonomic networks that support inhibitory, goal-directed, high-order brain functions. The tenth cranial vagus nerve communicates peripheral information to and from the brainstem, with afferents projecting to higher-order, cognitive areas such as prefrontal cortex, anterior cingulate, and amygdala. Additionally, descending projections from the PFC to the brainstem and hypothalamic structures allow for bi-directional communication between the central nervous system and the ANS through the vagus nerve [10]. As such, high levels of vagally-mediated HRV are associated with superior executive function [30], working memory [11], and emotional regulation [31]. Cognitive training including working memory has demonstrated that vagal activity reflects enhanced cognitive control of prefrontal networks [32]. Although sleep is not typically measured across the cognitive training interventions, the current findings suggest that executive function improvement may be mediated by the strengthening of prefrontal-autonomic networks during sleep.

Parasympathetic vagal activity is highest during SWS compared to all other states of consciousness [33]. Vagal activity is strongly coupled with delta activity (< 4Hz) during SWS and vagal enhancement precedes the onset of SWS [34]. Several studies have linked SOs with WM improvement. For example, studies have shown that fronto-parietal SOs, but not sigma, predicts WM improvement [16, 35]. However, not all studies report a consistent association between SOs and WM [13, 36, 37], and few accounts for autonomic activity. Chen and colleagues reported that vagal activity during SWS was a better predictor of WM improvement than SWA or vagal activity during wake [19]. In the current work, we found that changes in vagal autonomic activity during SWS, but not SOs per se, was critical for WM performance improvement. This, together with prior findings, suggests a non-negligible role of vagal influence on WM plasticity.

Given that both LTM and WM appear to rely on NREM sleep, one clear question emerges: How are the limited resources of NREM sleep shared across cognitive processes? The current findings are consistent with the hypothesis that competitive neural dynamics during NREM sleep underlie cognitive improvement. Supporting this hypothesis, prior research has shown that vagal nerve stimulation activates neurons in the locus coeruleus (LC) and increases NE levels in the brain [38, 39], and inactivation of LC impairs WM acquisition, while having no effect on consolidation or retention of spatial memories [40–44], whereas upregulating GABAergic networks impaired WM performance [45]. On the other hand, using ripple-triggered fMRI in monkeys, Logothetis and colleagues demonstrated that ripples orchestrate a privileged state of enhanced central brain activity by silencing output from the diencephalon, midbrain and brainstem, regions associated with autonomic regulation, which may serve to boost communication between hippocampus and cortex [20]. In addition, in both humans and mice, Lecci et al. (2017) demonstrated that heart rate and sigma power oscillate in antiphase with each other at 0.02 Hz, suggesting a periodic switch between sigma and autonomic activation every 50 seconds [46].

Here, using effective connectivity, we demonstrated that a GABAergic agonist enhanced naturally occurring cortical sigma dominance over vagal autonomic activity. Similar vagolytic findings have been shown with zolpidem in persistent vegetative state patients [47]. Furthermore, the magnitude of this central sigma influence on vagal activity predicted the trade-off between overnight LTM and WM improvement. Together with the previous literature, these finding suggest that sigma-dependent processes, including GABAergic hippocampal-thalamocortical networks, and vagal-dependent processes, including noradrenergic frontal-autonomic networks, may compete for sleep resources during NREM sleep. We hypothesize that the shared resource may be the SOs, which when coupled with ripple-nested sigma, promotes LTM and suppresses other processes, and when uncoupled, facilitates WM by enhancing prefrontal-autonomic networks. We further hypothesize that sigma may act as a gating mechanism that regulates SO resources for other processes, which would explain the mixed findings of SOs for WM improvement. Given that approximately 20% of slow oscillations during NREM are sigma-coupled [48], this leaves plenty of resources to be divided amongst other processes, including WM.

These data suggest a trade-off in which the two memory processes (LTM and WM) alternate during NREM sleep via a complex interaction at the synaptic (GABA vs NE activation), systems (thalamocortical vs frontal-midbrain), and mechanistic level (sigma-coupled SO vs uncoupled SO) (see graphical model in Fig. 6). Further research enhancing vagal activity and suppressing sigma activity is needed to show a double dissociation and tease apart these competitive mechanisms. Future work is also required to test the generalizability across multiple cognitive domains (i.e. motor learning) and tasks (i.e. non-associative LTM and N-back WM tasks) that relies on NREM sleep. The sleep switch mechanism and separable sleep features associated with WM and LTM processing suggest directions for future translational research on cognitive disturbances observed in neurodegenerative disorders such as Alzheimer’s and Parkinson’s disease, both of which involve the decline of sleep [49, 50].

**Fig. 6.**
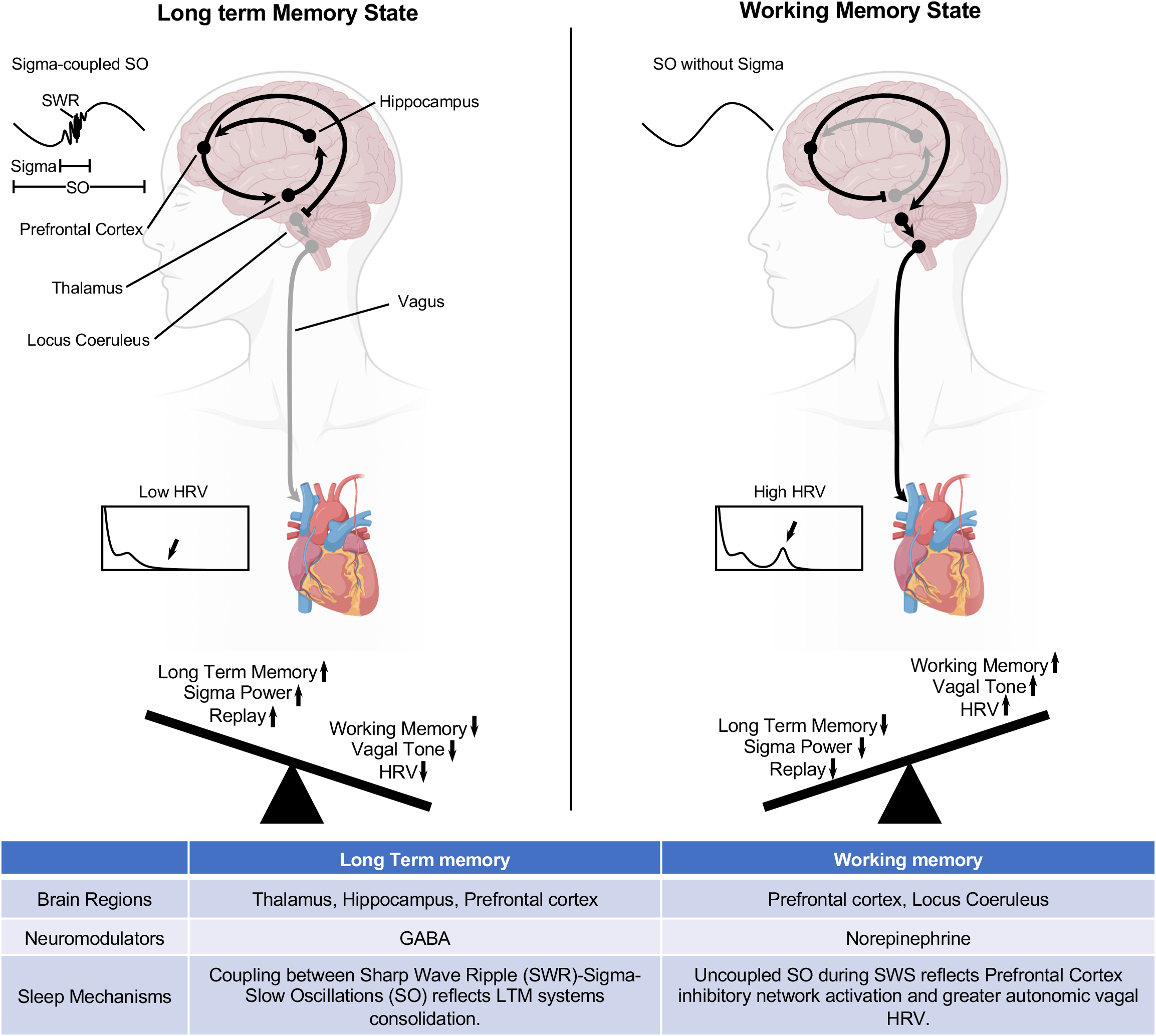
Sleep Switch Model. The model represents the proposed brain regions, primary neuromodulators, and sleep mechanisms involved in the Long-term memory state and the Working memory state that toggle throughout non-rapid eye movement (NREM) sleep. During the Long-Term Memory state, consolidation occurs via sigma-coupled SOs, which leads to reduced autonomic vagal-dependent activity and less WM improvement. During the Working Memory state, greater efficiency occurs during uncoupled SOs associated with increased autonomic vagal-dependent activity, which leads to reduced central sigma-dependent activity and less LTM consolidation.

### Limitations and future research

Limitations of this study include using a convenience sample of both men and women, and a lack of hormonal status among the young women, which can have an impact on cardiac vagal activity and sigma activity [52]. Future studies examining hormonal fluctuation are needed to understand the interaction between central sigma and ANS profiles during sleep and their impact on cognition. Additionally, though we did not measure respiration directly, we did analyze the frequency peak of HF (HFfp) in order to control for respiratory rate, which can affect the HRV. HFfp showed no difference between the two drug conditions and varied within a narrow range in the HF spectrum, between 0.22 and 0.26 Hz. Thus, it is unlikely that respiratory activity played a key role in zolpidem’s modulation on HRV and memory. However, we cannot completely exclude the effect of drug on cardiopulmonary coupling, as may be detected using measures of coherence. In addition, our experimental design did not include an adaptation night, and thus may have caused the “first-night effect”. However, the visits were counterbalanced by drug conditions, therefore the first-night effect should have canceled out across subjects. Furthermore, given that zolpidem is commonly prescribed to insomniacs, studies are needed to investigate if chronic use of zolpidem leads to WM deficits or biased memory trade-off during sleep. Lastly, due to methodological differences between EEG and ECG analyses, we measured sigma power as a proxy of spindles, which was not directly correlated with sleep-dependent behavioral changes. In addition, our study was limited by adhering to standard measures of vagal activity that require 5 min epochs, which reduced temporal specificity. This limitation constrains our effective connectivity analysis to all sleep epochs. We reported REM and wake free results in the supplementary (Figure S9, S10), which were in the same direction as what we reported in the main paper, but less robust, likely due to under power (Table S13). It’s therefore crucial that future research develop validated markers of vagal activity in shorter windows. Our results are lack of temporal specificity of sleep micro events and thus future research with a greater temporal precision around physiological events is needed to provide insight into shifts between central- and autonomic-dependent activities.

## Materials and Methods

### Participants

34 adults in experiment 1 (*M*_age_ = 20.88 ± 1.88 years, 17 Females) and 38 adults in experiment 2 (*M*_age_ = 20.85 ± 2.97 years, 19 Females) with no history of neurological, psychological, or other chronic illnesses were recruited for the study (*Supplemental Table S1* demographics). All participants signed informed consent, which was approved by the Western Institutional Review Board and the University of California, Riverside Human Research Review Board. Exclusion criteria included irregular sleep/wake cycles; sleep disorder; personal or familial history of diagnosed psychopathology; substance abuse/dependence; loss of consciousness greater than 2 minutes or a history of epilepsy; current use of psychotropic medications; and any cardiac or respiratory illness that may affect cerebral metabolism, which was determined during an in-person psychiatric assessment with trained research personnel. Additionally, all participants underwent a medical history and physical appointment with a staff physician to ensure their physical well-being. All subjects were naive to or had limited contact with (<2 lifetime use and no use in last year) the medication used in the study. Participants were asked to refrain from consuming caffeine, alcohol, and all stimulants for 24 h prior to and including the study day. Participants filled out sleep diaries for one week prior to each experiment and wore wrist-based activity monitors the night before the study (Actiwatch Spectrum, Philips Respironics, Bend, OR, USA) to ensure participants were well-rested (at least 7 hours per night during the week including the eve of the experimental day). Participants received monetary compensation and/or course credit for participating in the study. Study procedures were illustrated in Fig. 1.

### Data Reduction

#### Experiment 1

25 participants completed 4 visits (2 placebo nights and 2 zolpidem nights), 8 participants completed 2 visits (1 placebo night and 1 zolpidem night), 1 participant completed a zolpidem visit, due to scheduling conflicts. Therefore, 56 placebo and 59 zolpidem nights were included in the analyses.

#### Experiment 2

36 participants completed the placebo night, and 35 participants completed the zolpidem night PSG recordings. 35 participants completed all three sessions of operation-span (working memory) task in both placebo and zolpidem conditions. 33 participants completed all three sessions of word-paired associates (long-term memory) task in both placebo and zolpidem conditions.

### Sleep Recording

EEG data were acquired using a 32-channel cap (EASEYCAP GmbH) with Ag/AgCI electrodes placed according to the international 10-20 System (Jasper, 1958). 22 electrodes were scalp recordings and the remaining electrodes were used for electrocardiogram (ECG), electromyogram (EMG), electrooculogram (EOG), ground, an online common reference channel (at FCz location, retained after re-referencing), and mastoid (A1 & A2) recordings. The EEG was recorded with a 1000 Hz sampling rate and was re-referenced to the contralateral mastoid (A1 & A2) post-recording. Data were pre-processed using BrainVision Analyzer 2.0 (BrainProducts, Munich Germany). Eight scalp electrodes (F3, F4, C3, C4, P3, P4, O1, O2), the EMG, and EOG were used in the scoring of the nighttime sleep data. High pass filters were set at .3 Hz and low pass filters at 35 Hz for EEG and EOG. Raw data were visually scored in 30-sec epochs into Wake, Stage 1, Stage 2, Slow Wave Sleep (SWS) and rapid eye movement (REM) sleep according to the Rechtschaffen & Kales’ manual using HUME, a custom MATLAB toolbox. After staging, all epochs with artifacts and arousals were identified rejected by visual inspection before spectral analyses. Minutes in each sleep stage and sleep latencies (SL) (the number of minutes from lights out until the initial epoch of sleep, Stage 2, SWS and REM) were calculated. Additionally, wake after sleep onset (WASO) was calculated as total minutes awake after the initial epoch of sleep, and sleep efficiency (SE) was computed as total time spent asleep after lights out (~11:00PM) divided by the total time spent in bed (~11:00PM-9:00AM) * 100. Sleep architectures were reported in *Supplemental Table S2 and S3*.

### Heart Rate Variability

Electrocardiogram (ECG) data were acquired at a 1000-Hz sampling rate using a modified Lead II Einthoven configuration. We analyzed HRV of the R-waves series across the whole sleep/wake period using Kubios HRV Analysis Software 2.2 (Biosignal Analysis and Medical Imaging Group, University of Kuopio, Finland), according to the Task Force guidelines [57]. RR peaks were automatically detected by the Kubios software and visually examined by trained technicians. Incorrectly detected R-peaks were manually edited. Missing beats were corrected via cubic spline interpolation. Inter-beat intervals were computed, and a third-order polynomial filter was applied on the time series in order to remove trend components. Artifacts were removed using the automatic medium filter provided by the Kubios software.

The HRV analysis of the RR series was performed by using a Matlab-based algorithm. An autoregressive model (Model order set at 16) was employed to calculate the absolute spectral power (ms2) in the LF HRV (0.04–0.15 Hz; ms2) and the HF HRV (0.15–0.40 Hz; ms2; an index of vagal tone) frequency bands, as well as total power (TP; ms2; reflecting total HRV), and HF peak frequency (HFpf; Hz; reflecting respiratory rate). From these variables, we derived the HF normalized units (HF_nu_ = HF[ms^2^]/HF[ms^2^]+LF[ms^2^]) and the LF/HF ratio (LF[ms^2^]/HF[ms^2^]), an index often considered to reflect the sympathovagal balance (i.e., the balance between the two branches of the ANS), but whose meaning has been recently put into question. The LF, HF, and TP measures had skewed distributions and as such were transformed by taking the natural logarithm. Since the LF normalized units are mathematically reciprocal to HF_nu_ (i.e. LF_nu_ =1-HF_nu_), to avoid redundancy, only the HF_nu_ index is computed, an index often thought to reflect vagal modulation. Due to controversies about the physiological mechanisms that contribute to changes in LF activity, LF, LF/HF ratio and HFnu are difficult to make for these parameters, but they are reported for descriptive purposes.

In addition to the frequency domain parameters, RMSSD (ms; root mean square of successive differences) was calculated as a measure of vagally-mediated HRV in the time-domain. Similar to the frequency adjustments, to adjust for skewed distributions in the RMSSD, we report the natural logarithm. Additionally, RR (ms; time interval between consecutive R-peaks, reflecting frequency of myocardial contraction) were calculated as an index of cardiac autonomic control in our analyses.

For time-domain and frequency-domain HRV measures during different sleep stages, consecutive artifact-free 5-min windows of undisturbed sleep were selected across the whole night using the following rules: (a) the 1.5-min preceding, and (b) the entire 5-min epoch selected must be free from stage transitions, arousal, or movements. Windows were identified and averaged within Stage 2 sleep, slow-wave sleep (SWS), and REM sleep. We also analyzed 5 min of pre-sleep wakefulness (Rest). Epochs of N1 were not analyzed. All the HRV parameters by drug condition and sleep stage were reported in Supplemental Table S4-S6.

### Power spectral analysis

The EEG power spectrum was computed using the Fast Fourier Transformation. SWA (0.5-2Hz), delta (1-4Hz), theta (4-8Hz), alpha (8-13Hz), sigma (12-15Hz), beta (15-30Hz), and total power (0.3-35Hz) were calculated for each sleep stage (Stage 2, SWS and REM). The EEG epochs that were contaminated by muscle and/or other artifacts were rejected using a simple out-of-bounds test (with a ±200 μV threshold) on high-pass filtered (0.5 Hz) version of the EEG signals. Then, the normalized power spectra (% power of each frequency band of interest/ total power) were averaged bilaterally within each sleep condition/stage/subject. Power analyses that showed significant drug effect were reported in Supplemental Table S7 and S8.

### Effective Connectivity

To explore the causal information flow between CNS and ANS sleep features, we considered sigma to reflect CNS activity and HFln to reflect ANS activity. Sigma power of eight EEG channels (F3, F4, C3, C4, P3, P4, O1, O2) and HF of HRV were considered as signals to estimate effective connectivity. To adopt uniform timing across signals and avoid temporal misalignments between EEG signals and HF time series, a sliding window technique was incorporated with window length of 5 minutes and stride of 5 seconds. All data during nighttime sleep was used to have continuous time series of Sigma powers and HF, and length of 5 minutes was selected to be consist with HRV process. Therefore, for each subject, nine different signals were constructed including ratio of Sigma power band to total power of EEG of eight channels and HF power of HRV for each five-minute window (see Fig. 3a).

Generalized partial direct coherence (GPDC) measure was used to estimate causal information flow between Sigma power and HF. GPDC uses multivariate vector autoregressive (MVAR) model to model causal interactions between signals and estimate directed causal information flow between signals by using the coefficients and parameters of MVAR.

After constructing Sigma power and HF signals, GPDC was computed for each window with length of 500 samples (500 * 5 s = 2500 s) with stride of 250 samples. First, signals interactions were modeled by MVAR model (Eq. 1).

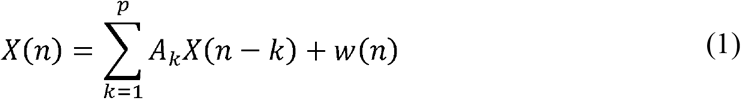

Where *X(n)* is the vector of signal values (with length of *N*, the number of signals, *N* = 8) in time *n*, *X(n)* = [*x*_1_(*n*), *x*_2_(*n*), …, *x_N_*(*n*)]^*T*^. *p* order of the MVAR model which was selected according to Akaike criterion (AIC), *p* = 4. *A_k_* is the matrix of MVAR coefficients and each element *a_ij_(k)*, stands how much *j*-th signal in time *n* - *k* affects *i*-th signal in time *n* and *w(n)* is the vector of model’s additive Gaussian noise with zero mean and covariance matrix Σ. After modeling the interaction of the signals, GPDC was computed using frequency domain of coefficients and covariance matrix as:

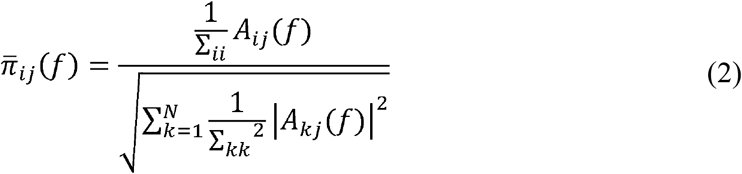

consequently:

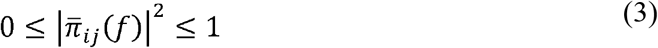

And

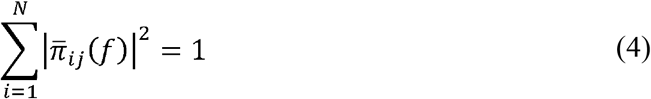

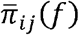 is the estimated matrix of causal information flow and the *j*-th column represent causal information outflow from the *j*-th signal to all the other signals. Average values over frequencies were considered for further process and based on the main purpose of the study two quantifier were defined as follow (see Fig. 3a):

1. Causal information outflow from HF to all EEG channels, HF_Outflow_ – Average (n=8) of causal information flow from HF to EEG sigma activity. HF_Outflow_ represents the strength of causal effect of HF to Sigma power.
2. Causal information inflow to HF from all EEG channels, HF_Inflow_ – Average (n=8) of causal information flow from EEG sigma activity to HF. HF_inflow_ represents the strength of causal effect of Sigma to HF.
3. Effective connectivity ratio: HFInflow over HFOutflow, where greater numbers represented a greater central sigma control over autonomic vagal activity than vice versa.

### Sigma/SO Coupling

Slow oscillations (SO) trough were detected for each channel automatically using the algorithm introduced by Dang-Vu et al. [58]. For each SO, the sigma power spectrum (12-16 Hz) was computed in the time margin of SO trough to 1s post SO trough. To access SOs which were coupled with Sigma waves, the median of all normalized Sigma power of SOs for all recording was computed for each channel. The SOs which had Sigma power greater than the median values in each quartile was considered as the SO-Sigma coupled and the number of coupled SOs was considered to further statistical analysis.

### Statistical Analyses

All statistical analyses were performed in R 3.6.2, using the libraries lme4 and lsmeans. P-values less than 0.05 were considered significant; p-values between 0.05 and 0.07 were considered trend-significant; p-values greater than 0.07 were considered non-significant. We used a linear mixed model (LMM) to evaluate the effects of zolpidem on sleep architecture, EEG power spectrum, autonomic profiles, and behavioral improvements. LMMs were chosen because it allows modeling of random effects and allow for the intercept and slope to be correlated [59]. LMMs are parametric models that use Maximum Likelihood Estimates (MLE) to obtain coefficients and covariance structures. LMMs do not depend on limited assumptions about variance-covariance matrix assumptions (sphericity). Additionally, LMMs allow inclusion of an unbalanced number of observations per participants in the analyses. Moreover, LMMs models take into account the influence of factors whose levels are extracted randomly from a population (i.e. participants), thus yielding more generalizable results.

#### Sleep architecture and Power spectrum

Using LMMs, we tested for the main effect of drug condition for sleep architecture (see Supplemental *Table S2 and S3),* EEG power spectrum (see Supplemental *Table S7 and S8)*.

#### Autonomic Profiles

For autonomic profiles, we tested for the main effect of drug condition and interactions between sleep stage and drug condition by approximating likelihood ratio tests (LRT) to compare LMMs with and without the effect of interest [60]. We first built a reduced (nested) model, with sleep stage as the only effect, and then included drug condition as a fixed effect in the full model. By comparing the reduced and full model using the LRT, we can interpret if drug condition significantly modulated the outcomes. Tukey’s correction for multiple testing was used for post-hoc comparisons.

#### Effective Connectivity

Using LMMs, we tested for the main effect of drug condition, the main effect of inflow vs outflow, and interaction between the two factors (see *Fig. 3b and 3c)*. We first built a reduced (nested) model, with inflow vs outflow, as the only effect, and then included drug condition as a fixed effect in the full model. By comparing the reduced and full model using the LRT, we can interpret if drug condition significantly modulated the outcomes. Tukey’s correction for multiple testing was used for post-hoc comparisons.

#### Behavioral Tasks

To investigate the drug effect on cognitive enhancement, LMMs were used with the drug condition as the predictor of interest (fixed effect), the improvement in WPA and OS tasks as outcome variables, and participants as crossed random effects. As we assume larger individual differences of improvement and difference in improvement between drug conditions, our LMMs include both a random intercept and a random slope term. To account for practicing effect on the tasks, we included visit and baseline performance as a covariate in the models. We first confirmed no differences at baseline (Test 1) between the placebo and zolpidem visits (*see Supplemental Fig. S1*). Next, we confirmed no differences of improvements across 12-hr of waking (Test 2 – Test 1) between the placebo and zolpidem visits (*see Supplemental Fig. S1*). We then tested the sleep-dependent changes in improvement: the overnight (Test 3 – Test 2) and 24-hr (Test 3 – Test 1) changes (see Fig. 4a and 4b). Again, we tested for the effect of drug condition by approximating LRTs.

#### Correlations

Lastly, we used a Pearson’s correlation coefficients to examine the functional roles of sigma, vagal activity, and causal information flow on sleep-dependent behavioral changes. We further used the Fisher r-to-z transformation to compare the differences between two correlations of interests.

