## Supplemental Figs. S1 to S10 Tables S1 to S13 for "Competitive dynamics underlie cognitive improvements during sleep"

**This PDF file includes:**

Figs. S1 to S10

Tables S1 to S13

Fig. S1. WPA and OS raw scores in Test1, Test2, and Test3, by Drug condition.

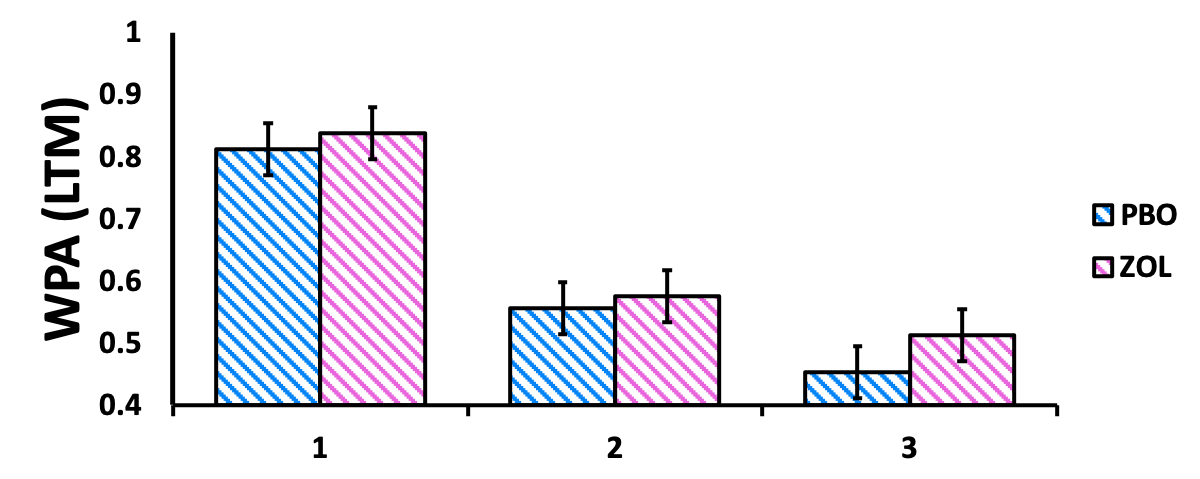

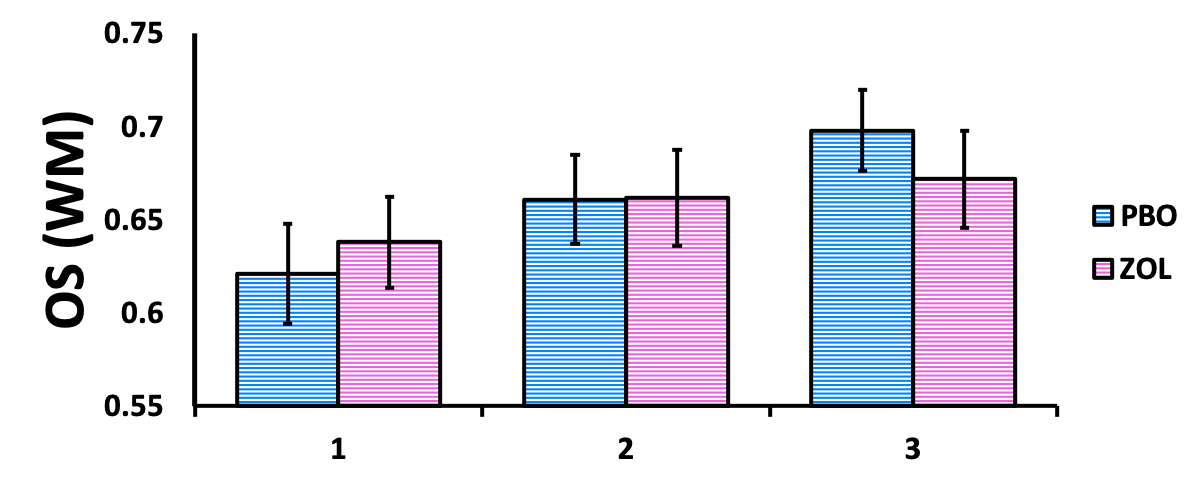

(a) For WPA, we estimated that the WPA Test 1 performance between the PBO vs ZOL visits are not significantly different (estimate= 0.0365, CI= (-0.1335, 0.2065), t= 0.4209, p= 0.6769), accounting for visit. The likelihood ratio test was not significant (LR = 1.4227; p = 0.4910), suggesting that adding the factor Drug did not significantly improved the model. In addition, the improvement of WPA across 12-hr of waking (Test 2 – Test 1) between the PBO vs ZOL visits are not significantly different (estimate= -0.0828, CI= (-0.3519, -0.0096), t= -0.8832, p= 0.3844), accounting for visit. The likelihood ratio test was not significant (LR = 3.3100; p = 0.5073), suggesting that adding the factor Drug did not significantly improve the model.

(b) For OS, we estimated that the OS Test 1 performance between the PBO vs ZOL visits are not significantly different (estimate= 0.0237, CI= (-0.1034, 0.1508), t= 0.3660, p= 0.7167), accounting for visit. The likelihood ratio test was not significant (LR = 1.7795; p = 0.7762), suggesting that adding the factor Drug did not significantly improve the model. In addition, the improvement of OS across 12-hr of waking (Test 2 – Test 1) between the PBO vs ZOL visits are not significantly different (estimate= 0.0413, CI= (-0.0684, 0.1511), t= 0.7385, p= 0.4657), accounting for visit. The likelihood ratio test was not significant (LR = 2.2089; p = 0.6974), suggesting that adding the factor Drug did not significantly improved the model.

Fig. S2. Experiment 1 HRV Profiles by Sleep Stage and Drug Condition (Whole Night)

(c)

(a)

(b)

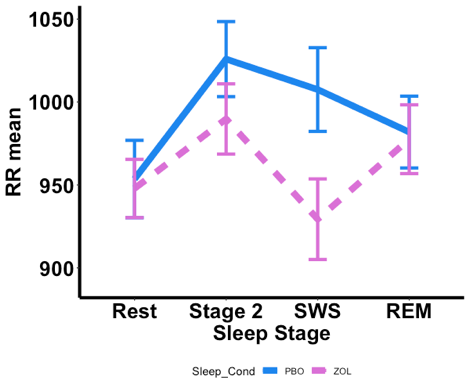

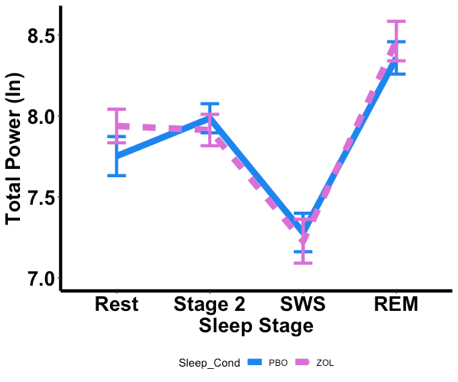

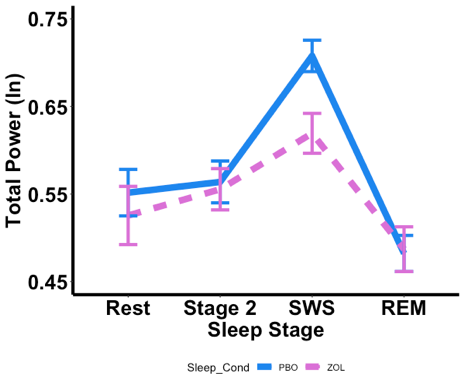

**HFnu**

(a) For RR mean, we report a significant effect of drug condition (F(1, 366) = 24.3443, p < .0001), with shorter heart rate intervals during the zolpidem condition; and a significant main effect of sleep stage (F(3, 366) = 14.2475, p < .0001), with slower heart rates during Stage 2 compared to Rest (p < .0001), REM (p = .0097), and SWS (p = .0001). We also found an interaction (F(3, 366) = 7.9875, p < .0001) between sleep stage and drug condition, with decreased vagal activity during SWS (p < .0001) and Stage 2 (p = .0041) in zolpidem compared with placebo, but not during REM (p = .6895), or Rest (p = .7436). The likelihood ratio test was significant (LR = 46.2681; p < .0001), suggesting that zolpidem significantly modulated the heart rates.

(b) For Total power (ln), we report a significant effect of sleep stage (F(3, 366) = 96.4240, p < .0001), with a lower power during SWS, compared to Rest, Stage 2, and REM sleep (all ps < .0001). No main effect (p = .3532) or interaction (p = .1622) of drug were found. The likelihood ratio test was not significant (LR = 6.0976; p = .1920), suggesting that zolpidem did not significantly modulate the total power.

(c) HFnu, we report no significant main effect of drug condition (F(1, 366) = 7.5959, p = .0061), with a lower normalized HF power during the zolpidem condition; and a significant main effect of sleep stage (F(3, 366) = 71.2954, p < .0001), with a greater normalized HF power during SWS, compared to Rest, Stage 2, and REM (all ps < .0001), and a lower normalized HF power during REM compared to Rest (p = .0001) and Stage 2 (p < .0001). The likelihood ratio test was significant (LR = 23.5682; p < .0001), suggesting that zolpidem significantly modulated the normalized HF power.

Fig. S3. Experiment 2 HRV Profiles by Sleep Stage and Drug Condition (Q2 and Q3)

(c)

(b)

(a)

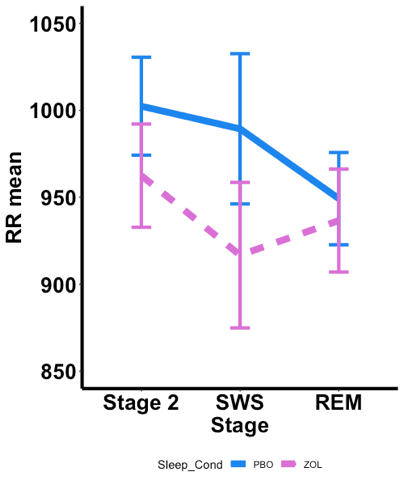

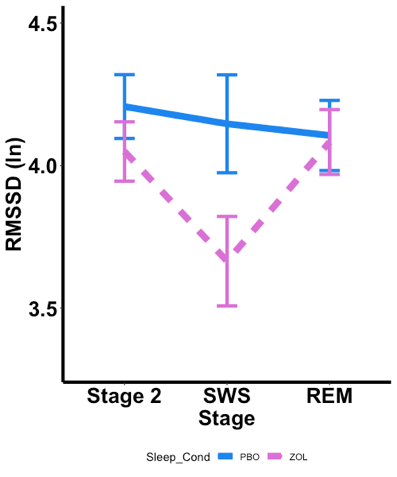

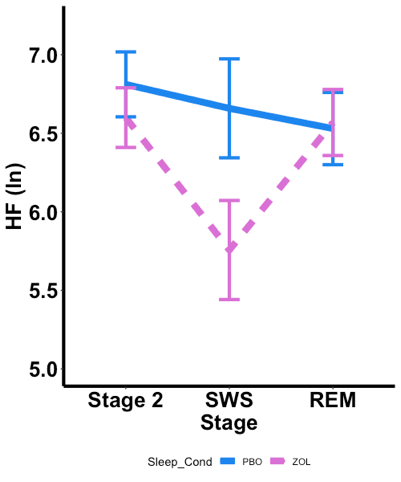

(d)

(e)

(f)

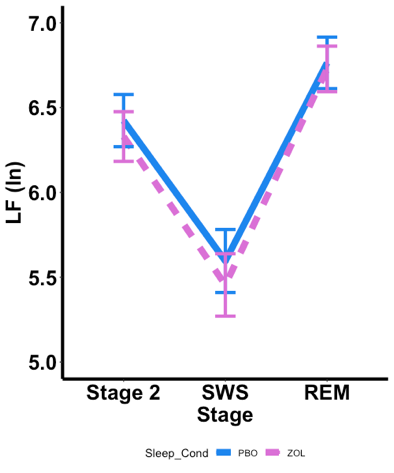

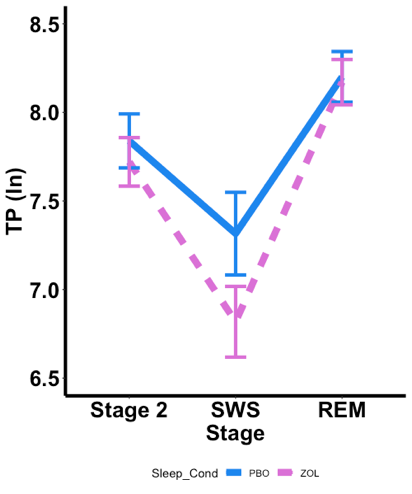

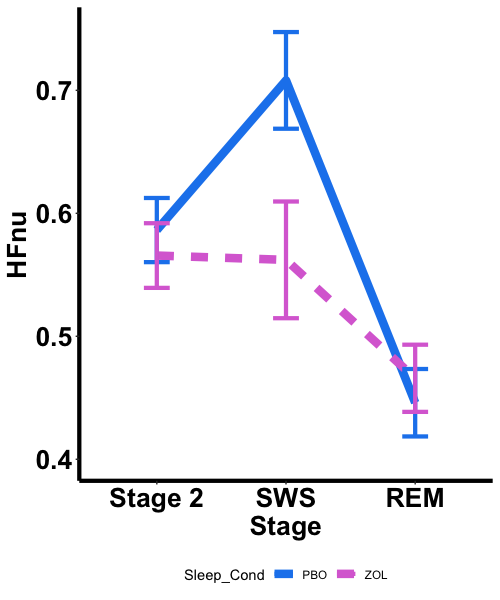

(a) For RR, we report a significant effect of Drug (F(1,153) = 12.2667, p = .0006), with shorter heart rate intervals during the Zolpidem condition; and a significant main effect of Stage (F(2,153) = 6.5458, p = .0019), with slower heart rate during Stage 2 compared to REM (p = .0016). No significant interaction (F(2,153) = 1.6530, p = 0.1949) was found. The likelihood ratio test was significant (LR = 15.3205; p = .0016), suggesting that adding the factor Drug significantly improved the model.

(b) For RMSSDln, we report a significant effect of Drug (F(1,153) = 11.1469, p = .0011), with lower vagal activity during the Zolpidem condition; and a significant main effect of Stage (F(2,153) = 7.9799, p = .0005), with SWS showing lower vagal activity compared to Stage 2 sleep (p = .0007) and REM (p = .0058). A significant interaction (F(2,153) = 3.6859, p = .0273) was found with decreased vagal activity during Stage 2 (p. = .0349) and SWS (p = .0002) but not REM (p = .7803) in Zolpidem compared with placebo. The likelihood ratio test was significant (LR = 18.1304; p < .0001), suggesting that adding the factor Drug significantly improved the model.

(c) For HFln, we report a significant effect of Drug (F(1,153) = 6.9588, p = .0092), with lower vagal activity during the Zolpidem condition; and a significant main effect of Stage (F(2,153) = 10.5279, p = .0001), with SWS showing lower vagal activity compared to Stage 2 sleep (p < .0001). A significant interaction (F(2,153) = 4.2946, p = .0153) was found with decreased vagal activity during SWS (p = .0004) but not Stage 2 (p = .1209) or REM (p = .8285) in zolpidem compared with placebo. The likelihood ratio test was significant (LR = 15.3518; p = .0015), suggesting that adding the factor Drug significantly improved the model.

(d) For LFln, we report a significant main effect of Stage (F(2,153) = 100.9869, p < .0001), with lower LF power during SWS compared to Stage 2 sleep (p < .0001) and REM. (p < .0001), as well as Stage 2 compared to REM (p < .0001). No significant effect of Drug (F(1,153) = 0.1914, p = .6623) or interaction (F(2,153) = 0.2151, p = .8067) was found. The likelihood ratio test was not significant (LR = 0.6396; p = .8873), suggesting that adding the factor Drug did not significantly improved the model.

(e) For TPln, we report a significant main effect of Stage (F(2,153) = 83.8500, p < .0001), with lower total power in SWS compared to Stage 2 sleep (p < .0001) and REM. (p < .0001), as well as Stage 2 sleep compared to REM (p < .0001). No significant effect of Drug (F(1,153) = 3.413, p = .0666) or interaction (F(2,153) = 1.5050, p = .2252) was found. The likelihood ratio test was not significant (LR = 6.5142; p = .0891), suggesting that adding the factor Drug did not significantly improved the model.

(f) For HFnu, we report a significant effect of Drug (F(1,153) = 6.9330, p = .0093), with lower vagal activity during the Zolpidem condition; and a significant main effect of Stage (F(2,153) = 65.2071, p < .0001), with higher vagal activity in SWS compared to Stage 2 sleep (p < .0001) and REM. (p < .0001), as well as Stage 2 sleep compared to REM (p < .0001). A significant interaction (F(2,153) = 8.3155, p = .0004) was found with lower vagal activity in the zolpidem condition compared with placebo during SWS (p < .0001) but not Stage 2 (p = .2466) or REM (p = .4754). The likelihood ratio test was significant (LR = 22.7421; p < .0001), suggesting that adding the factor Drug significantly improved the model.

Fig. S4. Experiment 2 HRV Profiles by Sleep Stage and Drug Condition (Whole Night)

(a)

(b)

(c)

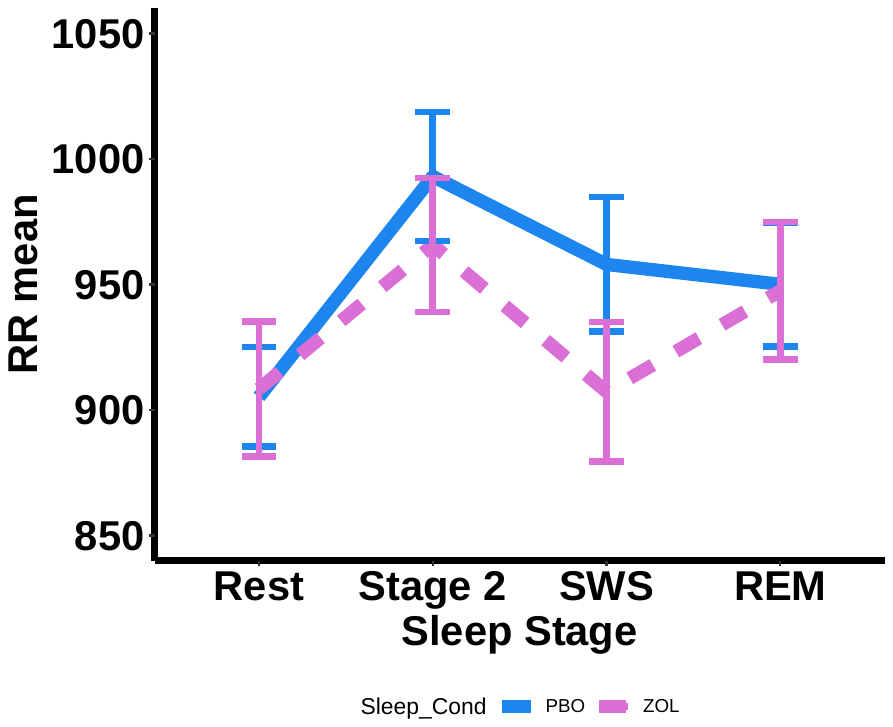

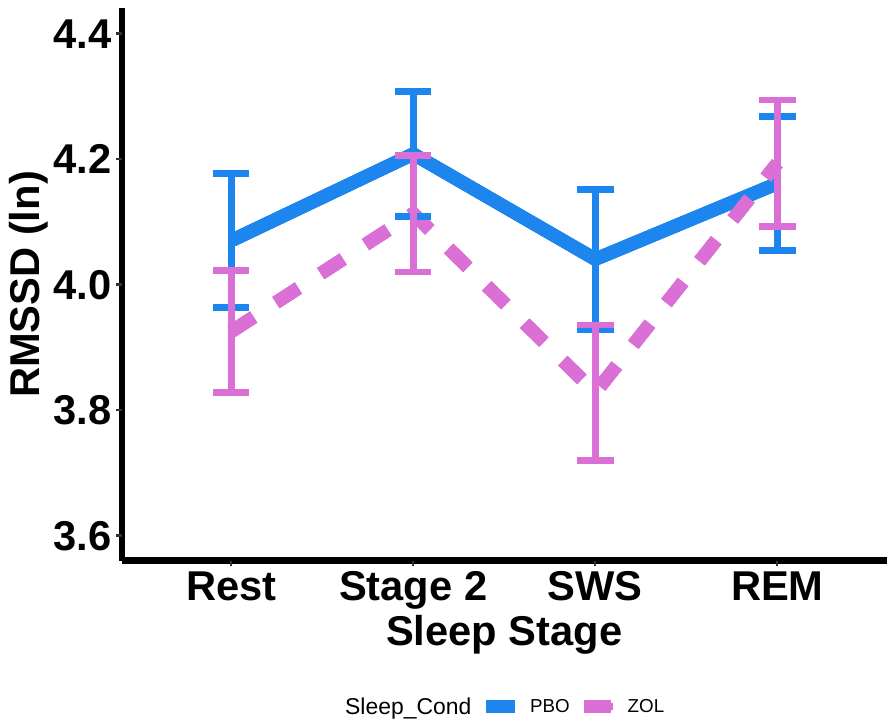

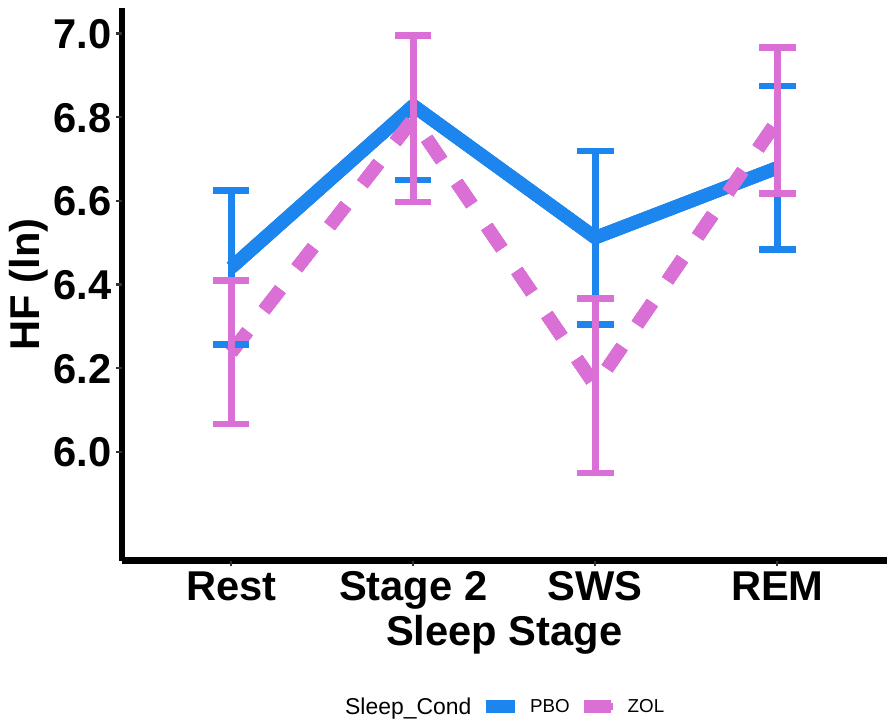

(d)

(e)

(f)

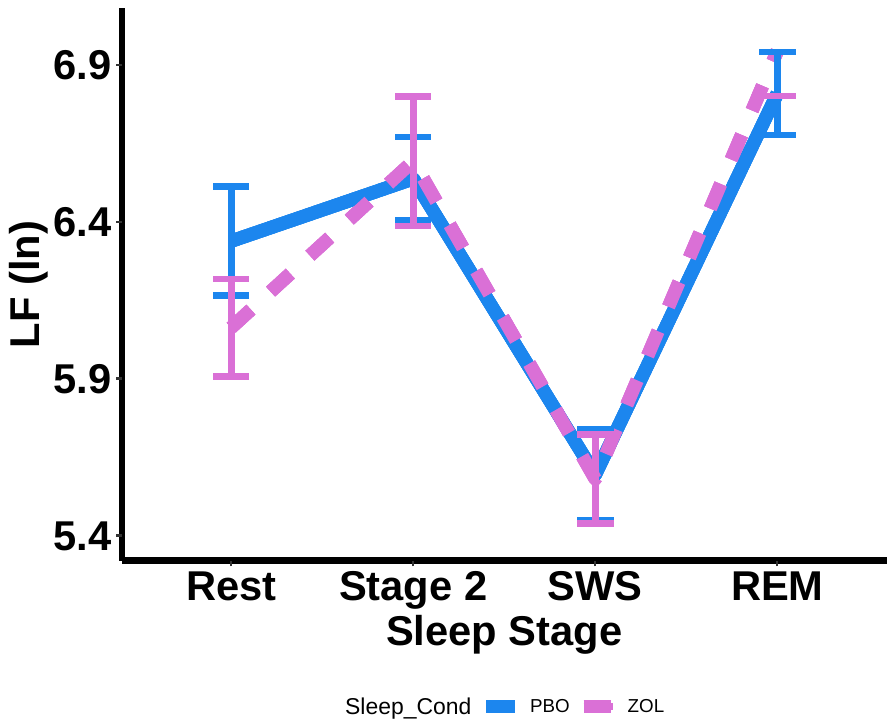

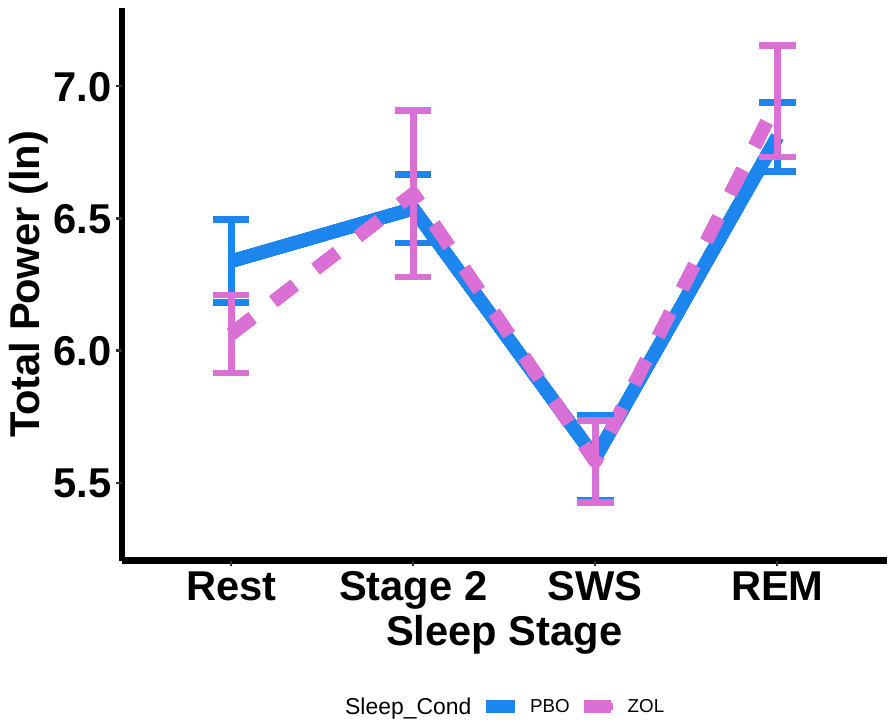

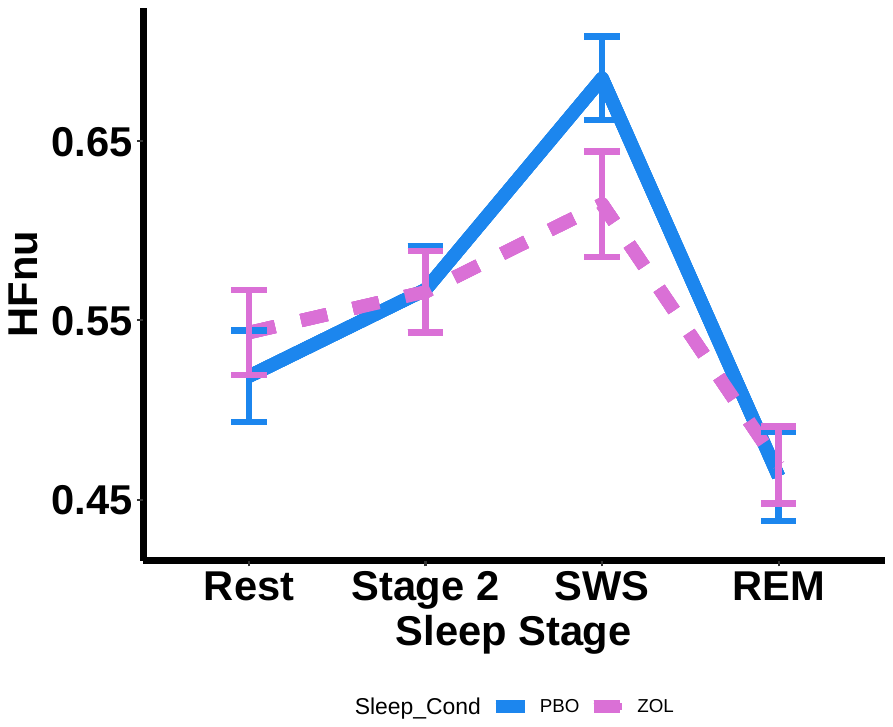

(a) For RR mean, we report a significant effect of drug condition (F(1, 222) =7.7956, p = .0057), with shorter heart rate intervals during the zolpidem condition; and a significant main effect of sleep stage (F(3, 222) = 10.2650, p < .0001), with slower heart rates during Stage 2 compared to Rest (p < .0001), REM (p = .0282), and SWS (p = .0005). We found no significant interaction (F(3, 222) = 1.6963, p = .1687) between sleep stage and drug condition. The likelihood ratio test was significant (LR = 12.9082; p = .0117), suggesting that zolpidem significantly modulated the heart rates.

(b) For RMSSDln, we report a significant effect of drug condition (F(1,222) = 4.1350, p = .0432), with lower vagal activity during the Zolpidem condition; and a significant main effect of Stage (F(3,222) = 10.8332, p < .0001), with SWS showing lower vagal activity compared to Stage 2 sleep (p < .0001) and REM (p < .0001). We found no significant interaction (F(3, 222) = 2.1063, p = .1003) between sleep stage and drug condition. The likelihood ratio test was significant (LR = 10.5405; p = .0322), suggesting that zolpidem significantly modulated RMSSD.

(c) For HFln, we report a significant effect of sleep stage (F(3,222) = 11.5488, p < .0001), with SWS showing lower vagal activity compared to Stage 2 sleep (p < .0001) and REM sleep (p = .0001), as well as a lower vagal activity during Rest compared to Stage 2 (p = .0024) and REM (p = .0297). No main effect of drug condition (p = .2949), or interaction (p = .0878) were found.

The likelihood ratio test was not significant (LR = 7.8393; p = .0976), suggesting that adding the factor Drug did not significantly improve the model. The effect of zolpidem on HFln was not as robust as experiment 1, likely due to under power, as the significant levels were very close to .05 and we have a lower sample size in experiment 2 compared to experiment 1.

(d) For LFln, we report a significant main effect of Stage (F(3,222) = 60.7116, p < .0001), with lower LF power during SWS compared to Stage 2 sleep (p < .0001) and REM (p < .0001), as well as Stage 2 compared to REM (p = .0098), Rest compared to Stage 2 (p = .0215), SWS (p < .0001) and REM (p < .0001). No significant effect of drug condition (F(1,222) = 0.3265, p = .5683) or interaction (F(3,222) = 0.8749, p = .4548) was found. The likelihood ratio test was not significant (LR = 3.0229; p = .5540), suggesting that adding the factor Drug did not significantly improve the model.

(e) For TPln, we report a significant main effect of Stage (F(3,222) = 30.7260, p < .0001), with lower total power in SWS compared to Stage 2 sleep (p < .0001) and REM (p < .0001), as well as Stage 2 sleep compared to REM (p = .0271). Total power during Rest was lower than Stage 2 (p = .0123) and REM (p < .0001), but greater than SWS (p = .0284). No significant effect of drug condition (F(1,222) = 0.646, p = .4226) or interaction (F(3,222) = 1.109, p = .3465) was found. The likelihood ratio test was not significant (LR = 4.0569; p = .3984), suggesting that adding the factor Drug did not significantly improve the model.

(f) For HFnu, we report a significant main effect of Stage (F(3,222) = 43.5761, p < .0001), with higher vagal activity in SWS compared to Stage 2 sleep (p < .0001), REM (p < .0001) and Rest (p < .0001), as well as Stage 2 sleep compared to REM (p < .0001). HFnu during REM was lower than Rest (p = .0001). No main effect of drug condition was observed (p = .2207). A significant interaction (F(3,222) = 2.9691, p = .0328) was found with lower vagal activity during SWS (p < .0001) but not Stage 2 (p = .2466) or REM (p = .4754) compared with placebo. The likelihood ratio test was significant (LR = 10.4879; p = .0330), suggesting that adding the factor Drug significantly improved the model.

Fig. S5. Experiment 2 HRV Profiles by Sleep Stage and Drug Condition (Q1)

(a)

(b)

(c)

(d)

(e)

(f)

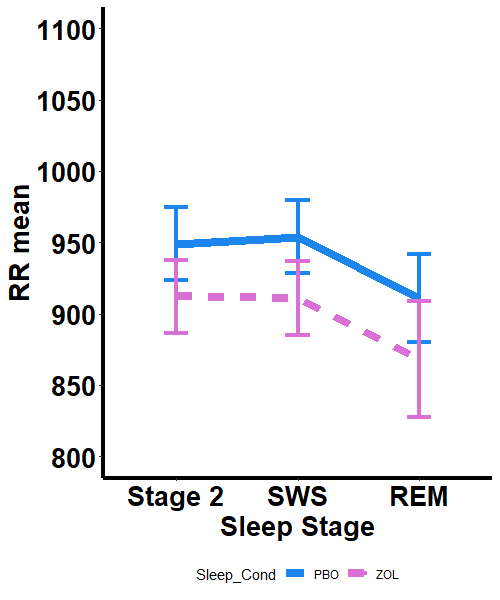

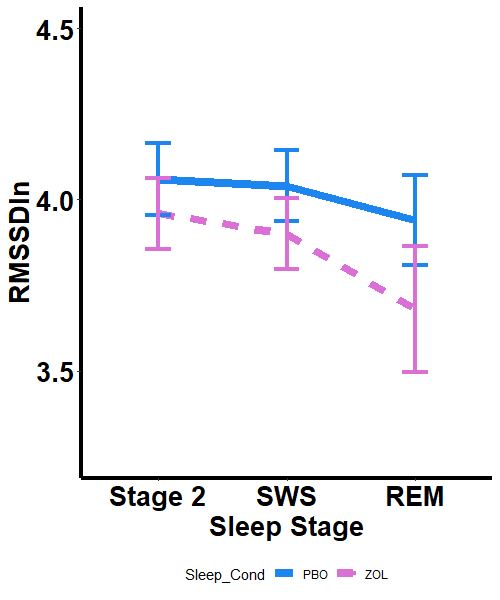

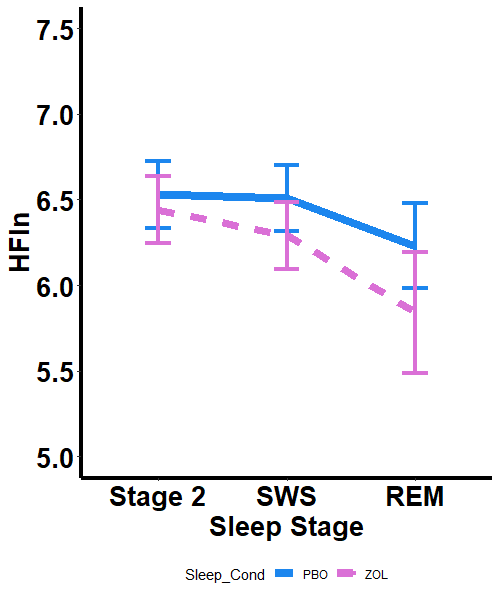

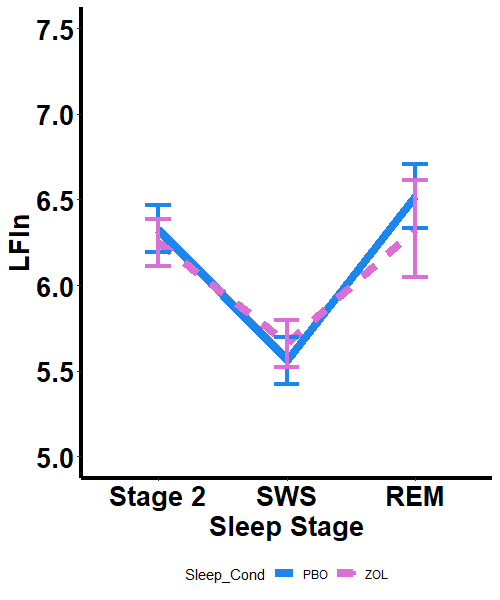

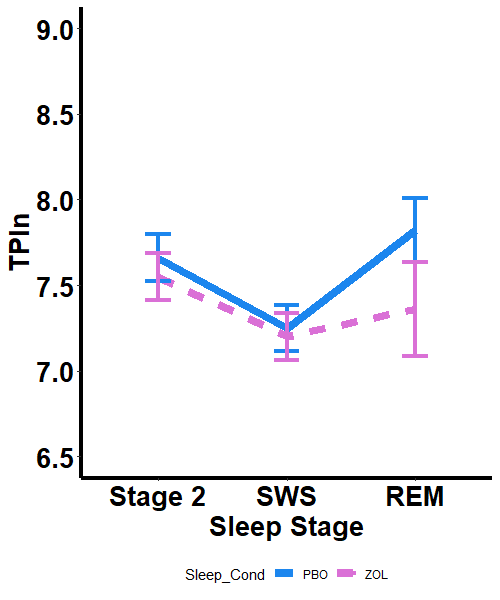

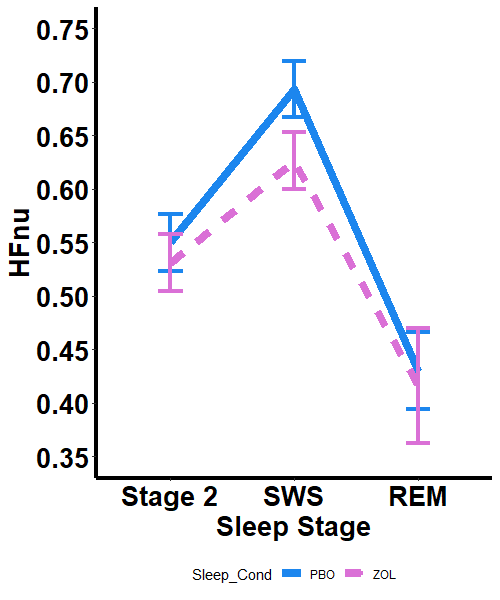

(a) For RR mean, we report a significant effect of drug condition (F(1, 108) = 15.9759, p = .0001), with shorter heart rate intervals during the zolpidem condition. No main effect of sleep stage (p = .1235) or interaction (p = .9591) were found. The likelihood ratio test was significant (LR = 15.5494; p = .0014), suggesting that zolpidem significantly modulatef the heart rates during quartile 1.

(b) For RMSSDln, we report a significant effect of drug condition (F(1, 108) = 8.0838, p = .0053), with lower vagal activity during the zolpidem condition. No main effect of sleep stage (p = .1002) or interaction (p = .7724) were found. The likelihood ratio test was significant (LR = 8.6199; p = .0348), suggesting that zolpidem significantly modulate RMSSD during quartile 1.

(c) For HFln, we report a significant effect of sleep stage (F(2,108) = 3.9421, p = .0223), with REM sleep showing lower vagal activity compared to Stage 2 sleep (p = .0078) and REM sleep (p = .0357). We found a significant effect of drug condition (F(1, 108) = 4.8879, p = .0292), with lower vagal activity during the zolpidem condition. No interaction between sleep stage and drug condition was found (p = .5593). The likelihood ratio test was not significant (LR = 6.1223; p = .1058), suggesting that zolpidem did not significantly modulate HF HRV during quartile 1.

(d) For LFln, we report a significant effect of sleep stage (F(2, 108) = 46.0026, p < .0001), with lower LF during SWS compared to Stage 2 (p < .0001) and REM (p < .0001). No main effect (p = .8326) or interaction (p = .4041) of drug were found. The likelihood ratio test was not significant (LR = 1.9326; p = .5865), suggesting that zolpidem did not significantly modulate LF HRV during quartile 1.

(e) For Total power (ln), we report a significant effect of sleep stage (F(2, 108) = 17.3386, p < .0001), with a lower power during SWS, compared to Stage 2 (p < .0001) and REM (p = .0276). No main effect (p = .1882) or interaction (p = .3786) of drug were found. The likelihood ratio test was not significant (LR = 3.7941; p = .2847), suggesting that zolpidem did not significantly modulate the total power during quartile 1.

(f) HFnu, we report a significant effect of sleep stage (F(2, 108) = 53.2369, p < .0001), with greater HFnu during SWS compared to Stage 2 (p < .0001) and REM (p < .0001), and greater HFnu during Stage 2 compared to REM (p = .0003). A main effect of drug condition was found (F(1, 108) = 5.5971, p = .0198). No interaction between drug condition and sleep stage (p = .3111) was found. The likelihood ratio test was significant (LR = 7.9951; p = .0461), suggesting that zolpidem significantly modulate the HFnu during quartile 4.

Fig. S6. Experiment 2 HRV Profiles by Sleep Stage and Drug Condition (Q4)

(a)

(b)

(c)

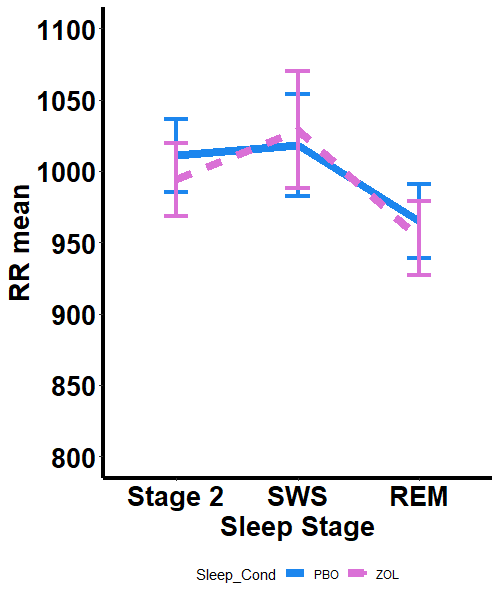

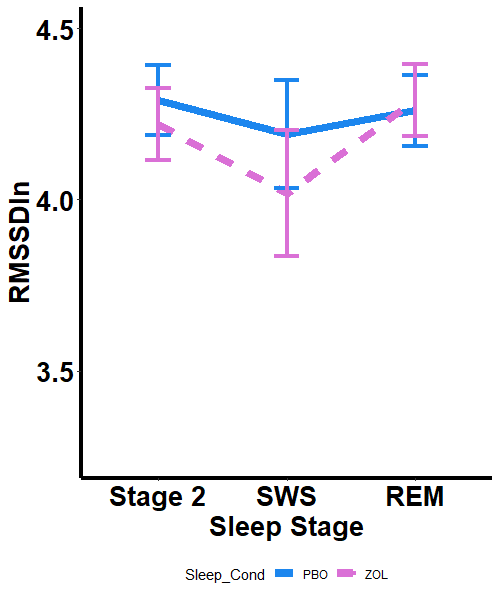

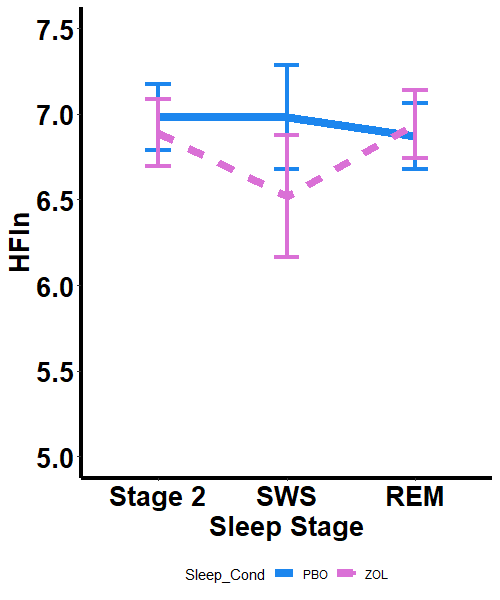

(d)

(e)

(f)

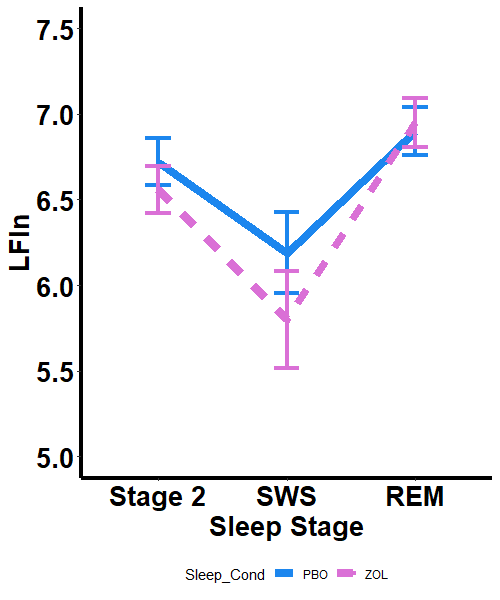

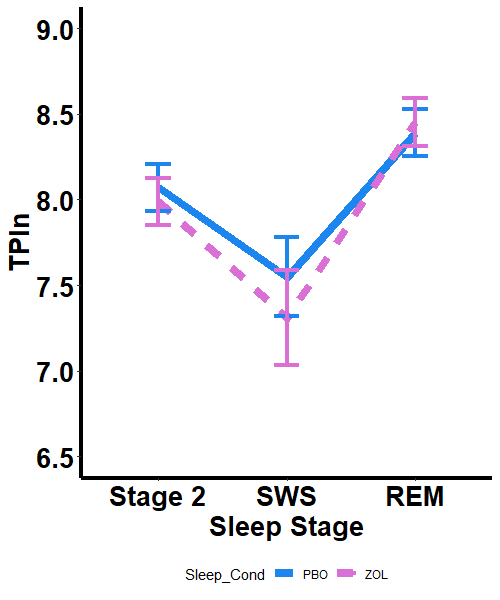

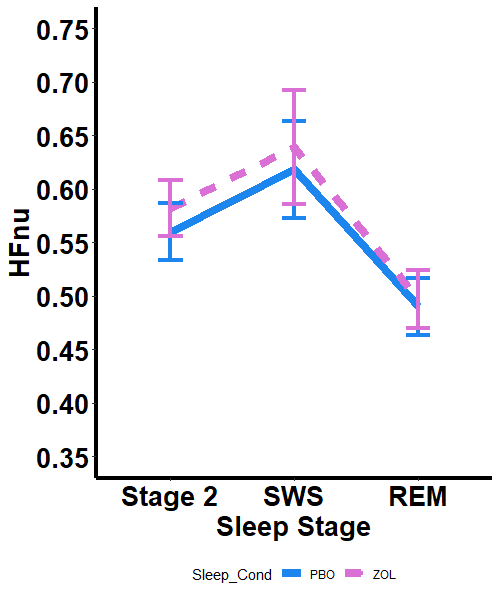

(a) For RR mean, we report a significant effect of sleep stage (F(2, 101) = 9.4608, p < .0001), with faster heart rates during REM compared to Stage 2 (p = .0003) and SWS (p = .0199). No main effect (p = .2798) or interaction (p = .3406) of drug were found. The likelihood ratio test was not significant (LR = 3.4397; p = .3287), suggesting that zolpidem did not significantly modulate the heart rates during quartile 4.

(b) For RMSSDln, no main effect of sleep stage (p = .4233), main effect of drug condition (p = .7777), or interaction (p = .4395) were found. The likelihood ratio test was not significant (LR = 1.7976; p = .6155), suggesting that zolpidem did not significantly modulate RMSSD during quartile 4.

(c) For HFln, no main effect of sleep stage (p = .5083), main effect of drug condition (p = .9742), or interaction (p = .5796) were found. The likelihood ratio test was not significant (LR = 1.3855; p = .7678), suggesting that zolpidem did not significantly modulate HF HRV during quartile 4.

(d) For LFln, we report a significant effect of sleep stage (F(2, 101) = 17.592, p < .0001), with greater LF during REM compared to Stage 2 (p = .0005) and SWS (p < .0001). No main effect (p = .5503) or interaction (p = .4056) of drug were found. The likelihood ratio test was not significant (LR = 2.2536; p = .5215), suggesting that zolpidem did not significantly modulate LF HRV during quartile 4.

(e) For Total power (ln), we report a significant effect of sleep stage (F(2, 101) = 25.925, p < .0001), with a greater power during REM, compared to Stage 2 (p < .0001) and SWS (p < .0001), as well as a greater power during Stage 2 compared to SWS (p = .0025). No main effect (p = .7794) or interaction (p = .6454) of drug were found. The likelihood ratio test was not significant (LR = .9959; p = .8022), suggesting that zolpidem did not significantly modulate the total power.

(f) HFnu, we report a significant effect of sleep stage (F(2, 101) = 15.9494, p < .0001), with lower HFnu during REM compared to Stage 2 (p < .0001) and SWS (p = .0003). No main effect (p = .3150) or interaction (p = .9229) of drug were found. The likelihood ratio test was not significant (LR = 1.2233; p = .7474), suggesting that zolpidem did not significantly modulate HFnu during quartile 4.

Fig. S7. Experiment 2 Effective Connectivity Modulated by Drug Condition

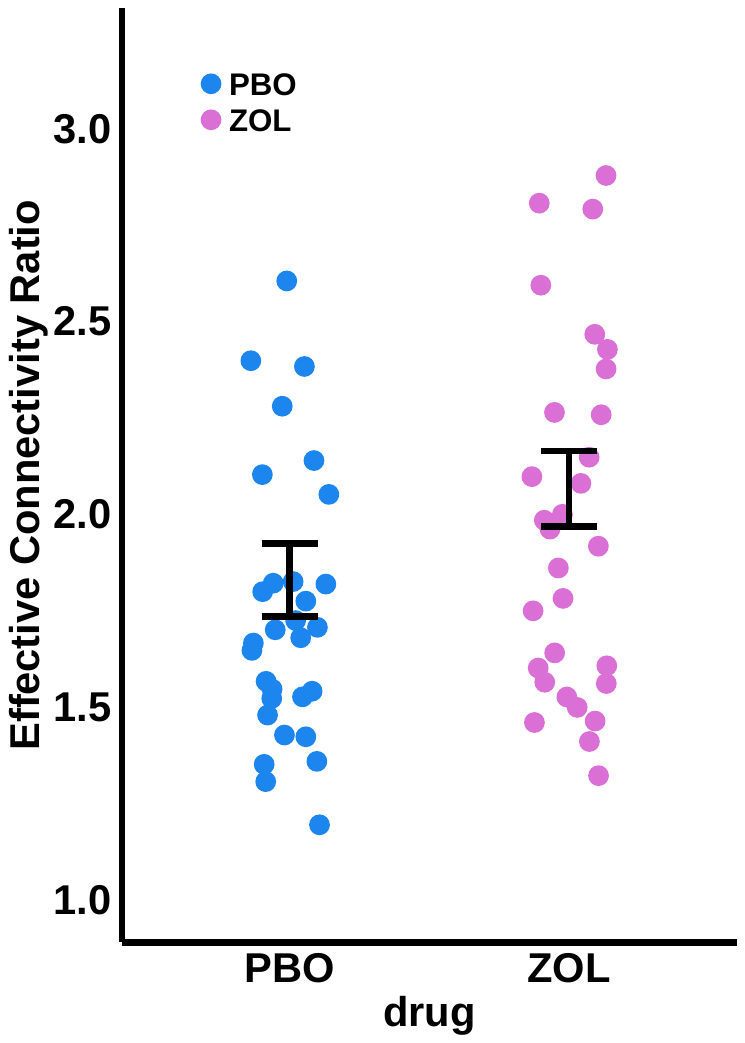

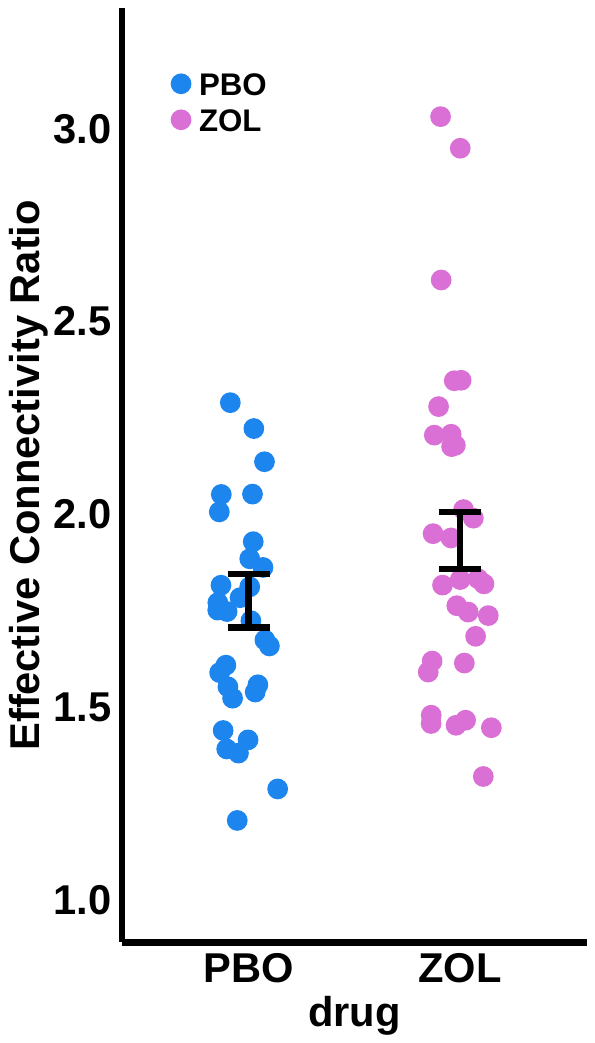

(a) Quartile 2 and 3 Effective connectivity ratios (HFInflow/ HFOutflow) increased significantly during the zolpidem condition (F(1, 32) = 5.4087, p = .0265).

(b) Whole-night Effective connectivity ratios (HFInflow/ HFOutflow) increased significantly during the zolpidem condition (F(1, 32) = 6.008, p = .0198).

Fig. S8 Experiment 2 Change in Spindle Density during SWS (Q2&3) correlated with Change in WPA Improvement between Drug Condition.

(c)

(b)

(a)

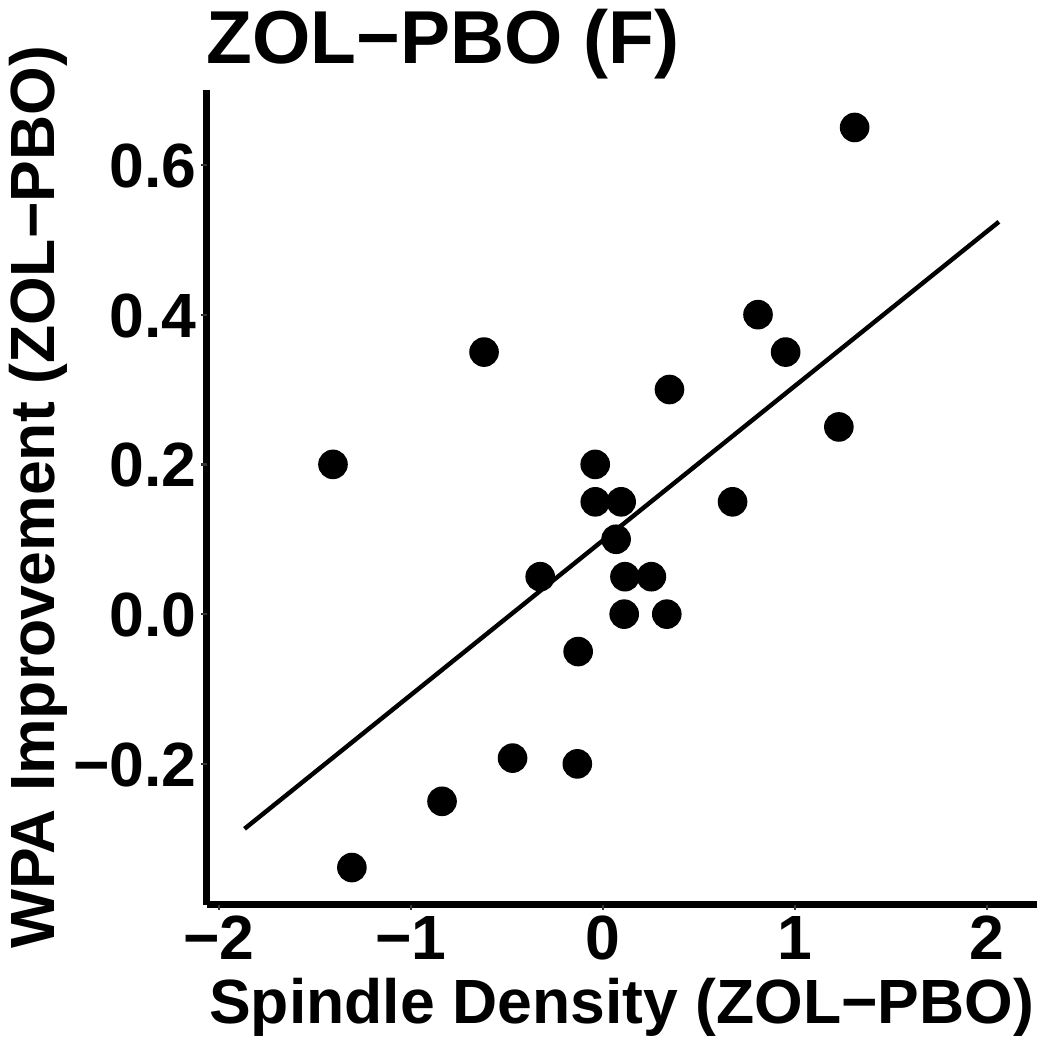

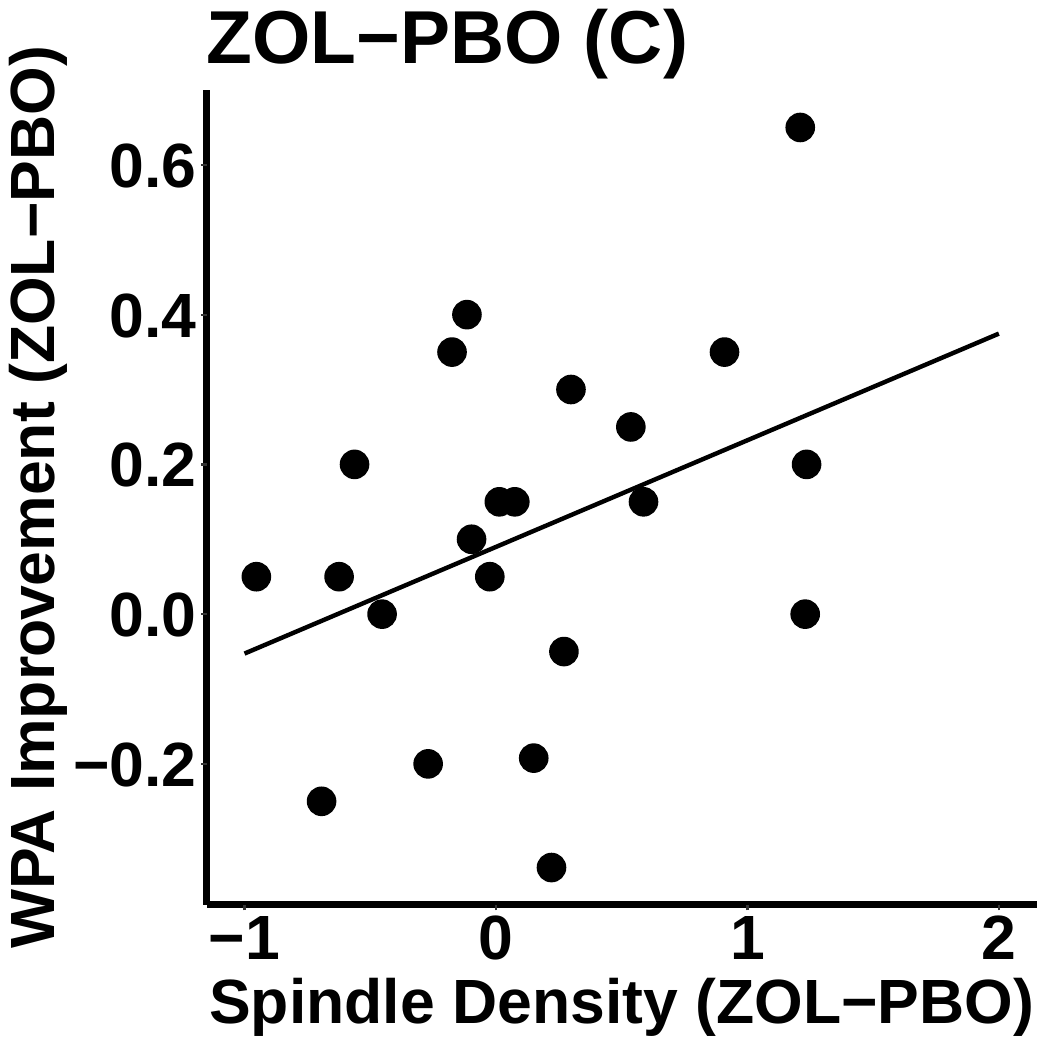

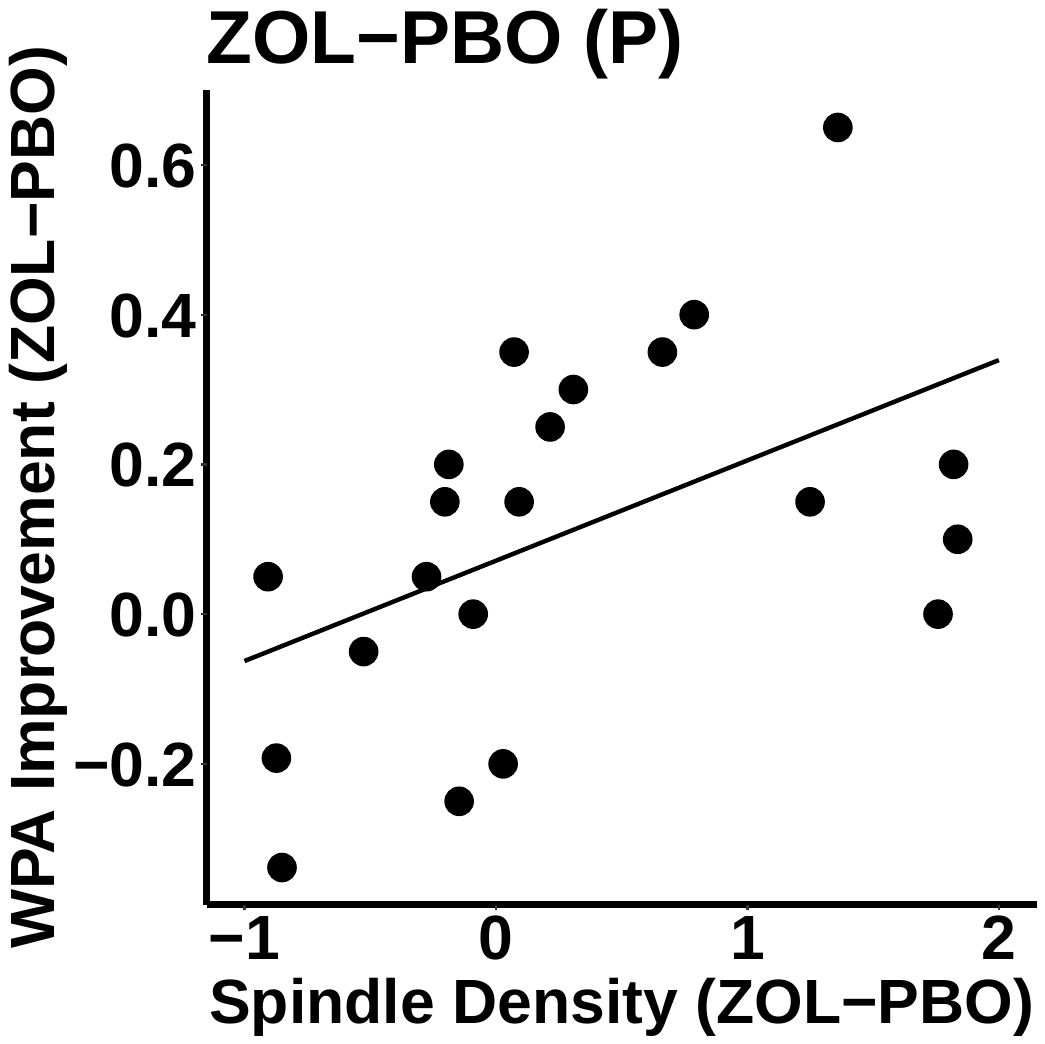

(f)

(e)

(d)

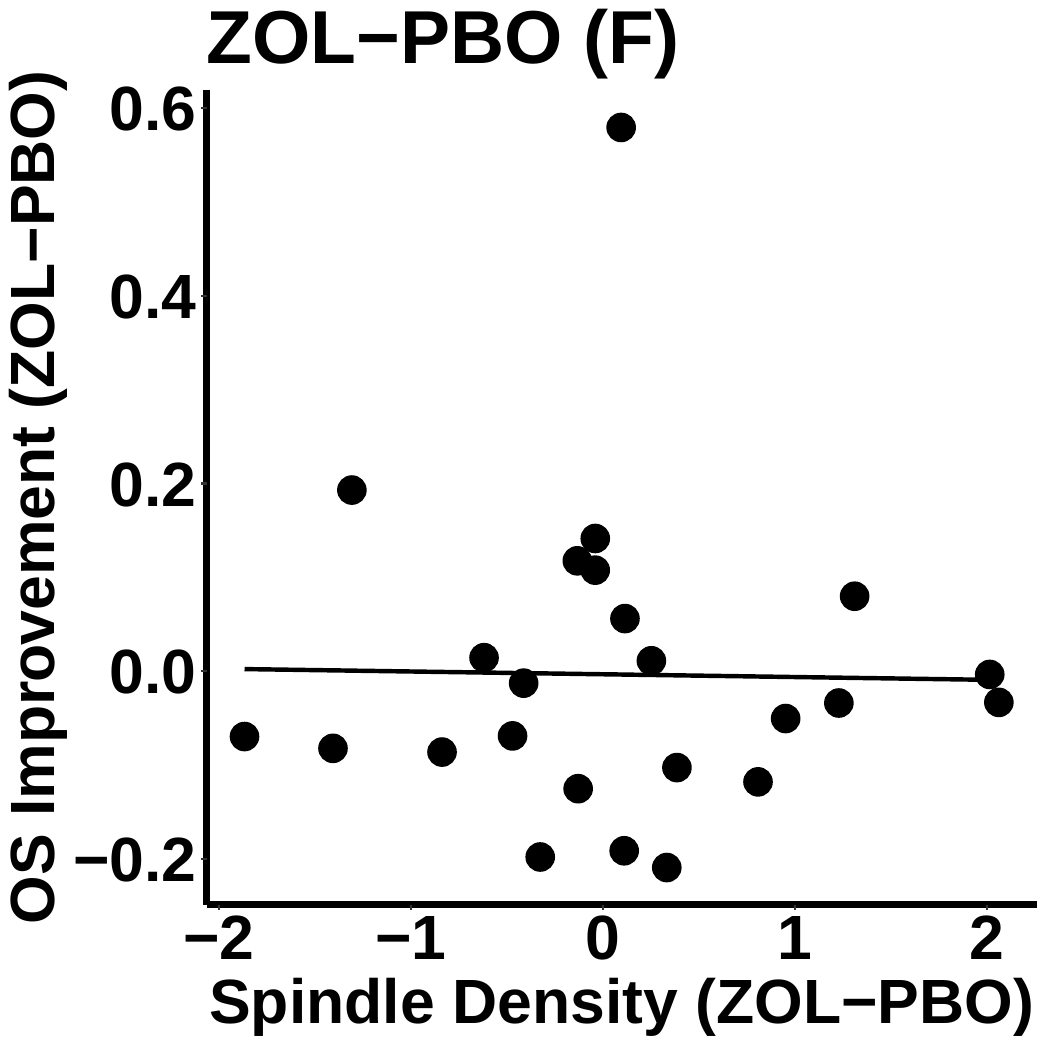

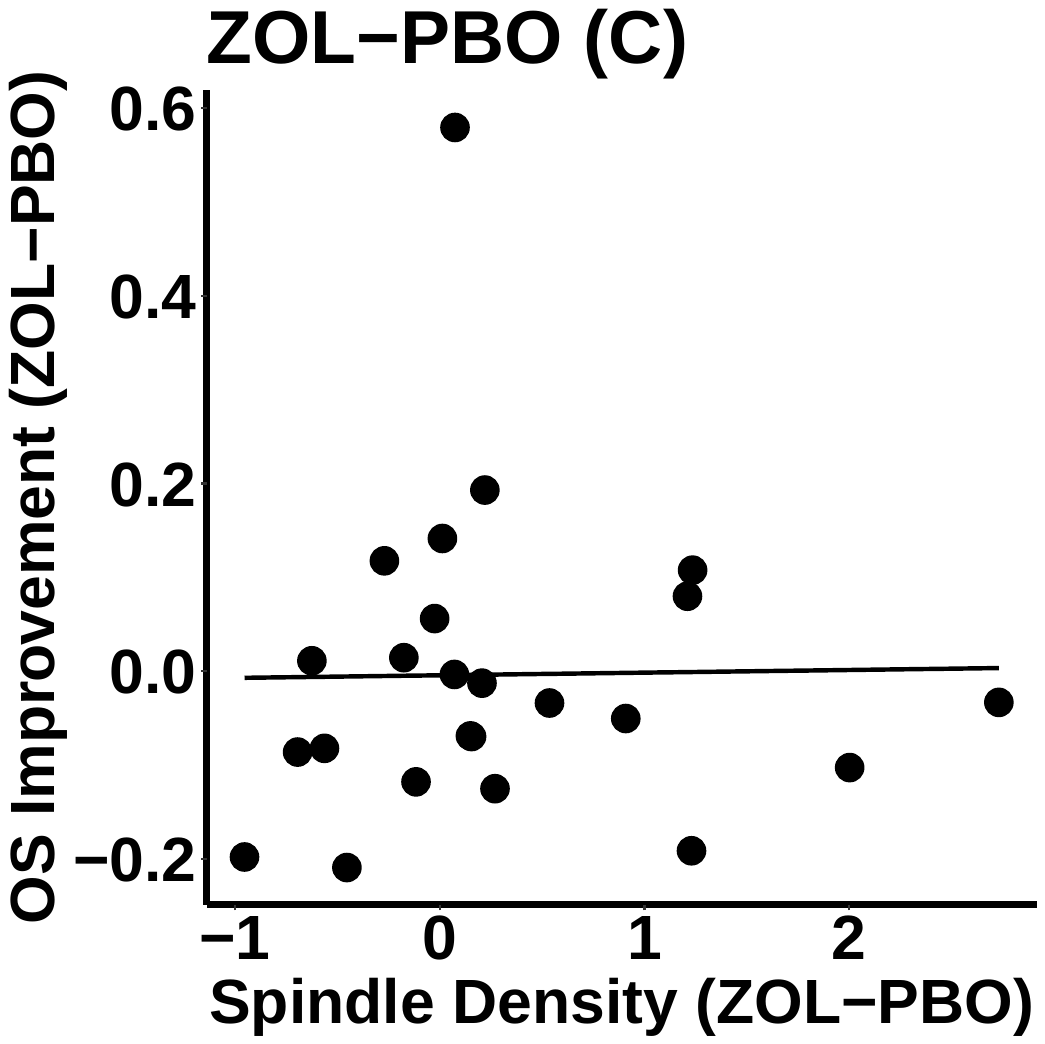

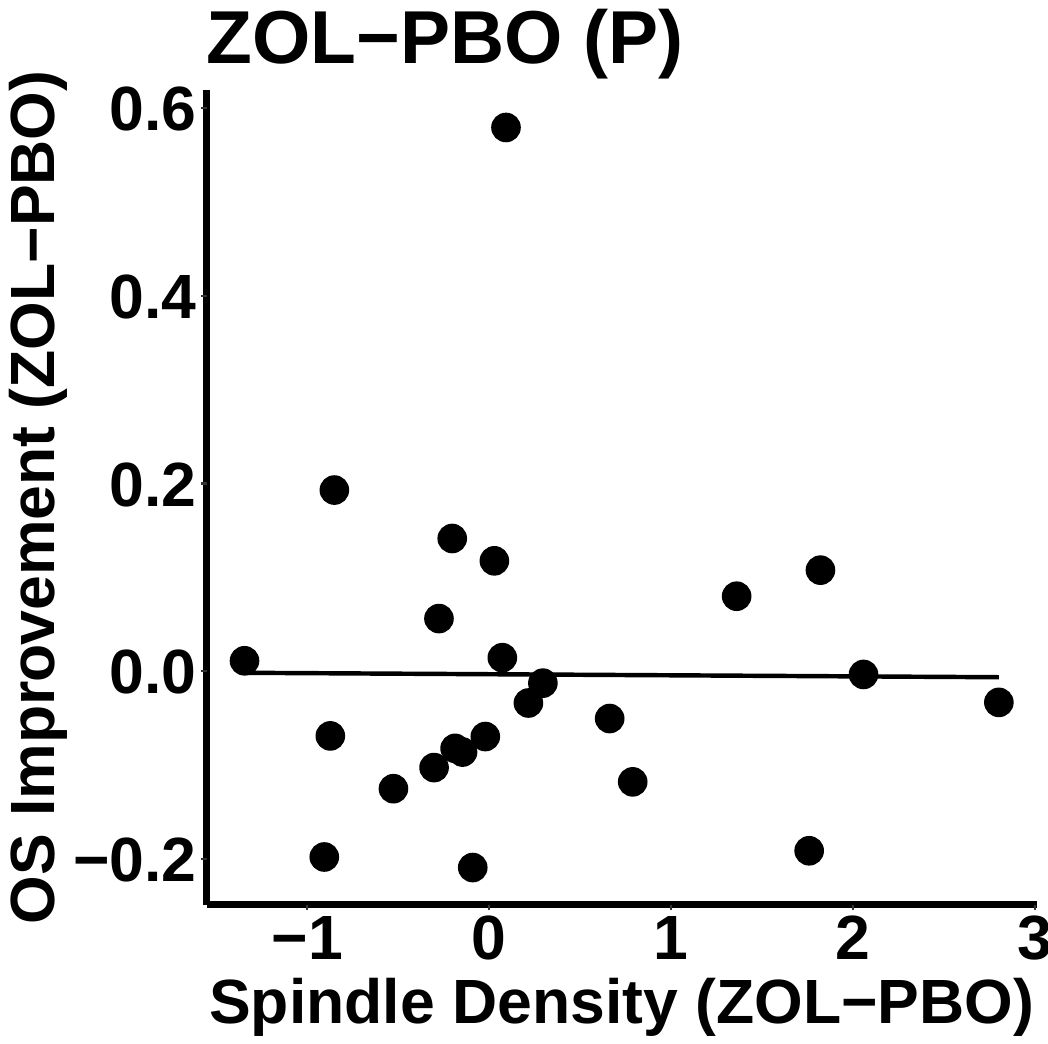

We observed a positive correlation between spindle density change (zolpidem minus placebo) and 24-hr LTM improvement (zolpidem minus placebo) in (a) frontal (r = .629; p = .0017) (b) central (r = .383, p = .0787), and (c) parietal (r = .475, p = .0254) regions. No such correlations were found in WM improvement in (d) frontal (r = -.018, p = .9346), (e) central (r = .015, p = .9443), or (f) parietal (r = -.007, p = .9733) regions. Channels were bilaterally averaged across F3, F4, C3, C4, P3, P4. For detail methods of spindle detection algorithm, please see Zhang et al. (2020) [7].

(c)

(b)

(a)

Fig. S9. Effective Connectivity Modulated by Drug Condition (NREM Only)

(a)

(b)

(c)

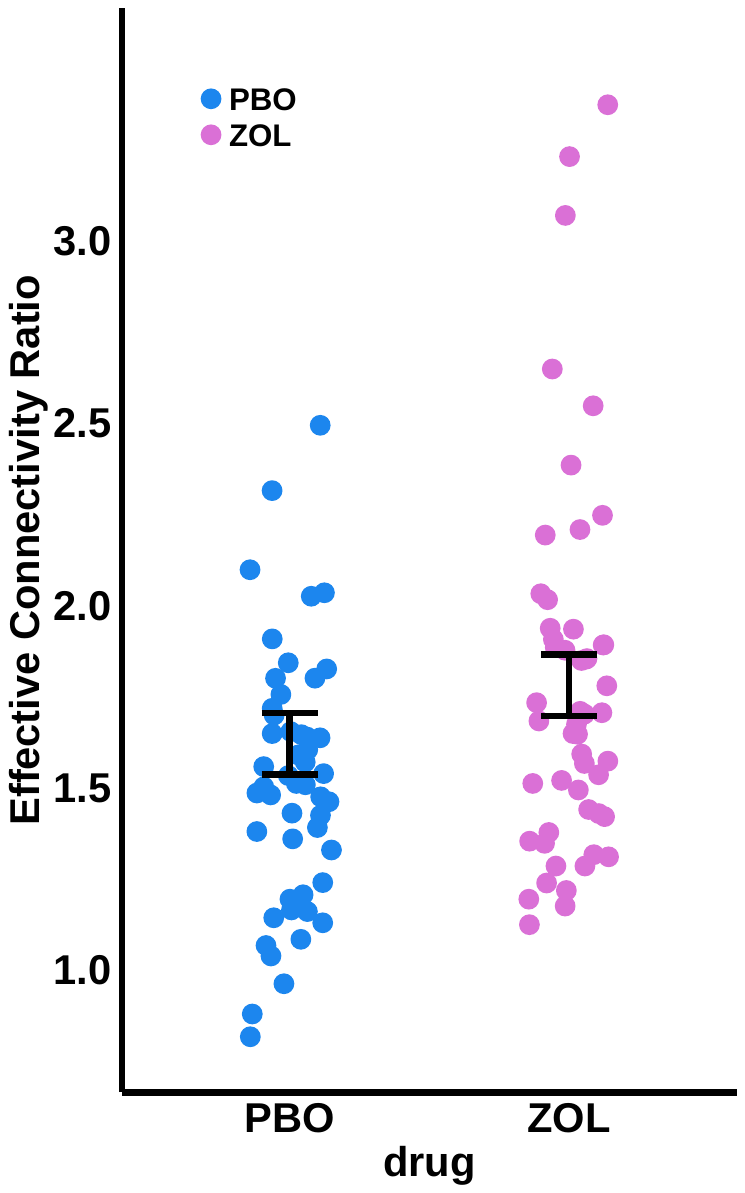

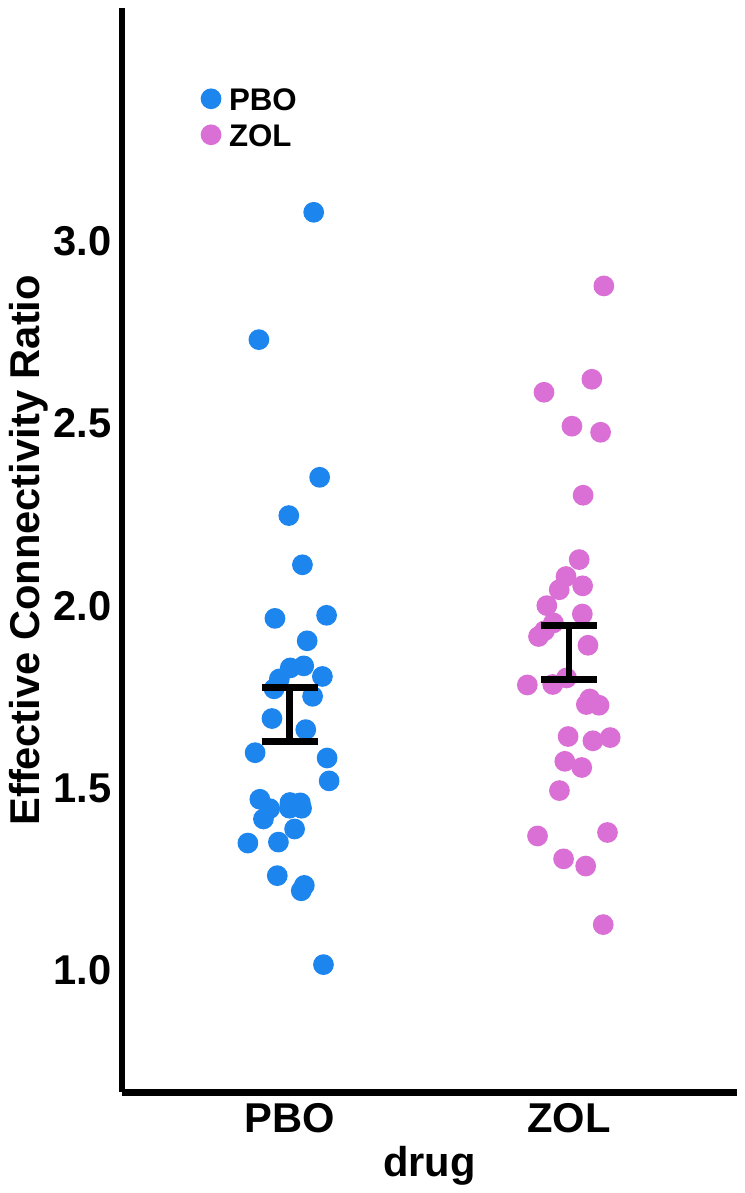

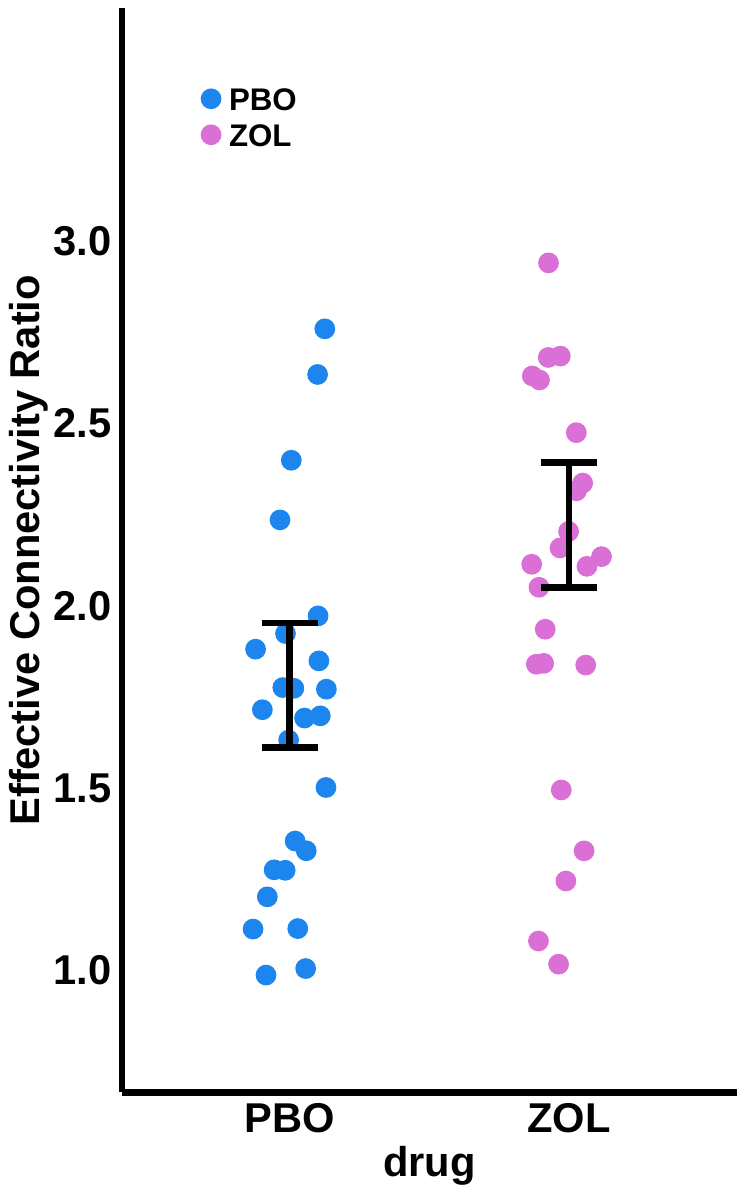

(a) In study 1, we found a tread difference in whole-night effective connectivity ratios (HFInflow/ HFOutflow) between drug conditions (F(1, 79) = 1.8252, p = .1810), with zolpidem increasing the effective connectivity ratio.

(b) In study 2, whole-night effective connectivity ratios (HFInflow/ HFOutflow) increased significantly during the zolpidem condition (F(1, 32) = 6.2615, p = .0176).

(c) In study 2, we next examined this drug effect in each quartile and found that quartile 3 fffective connectivity ratios increased during the zolpidem condition (F(1, 26) = 3.3029, p = .0807).

Fig. S10. Effective Connectivity Correlated with Behavioral Trade-off (NREM Only)

(a)

(b)

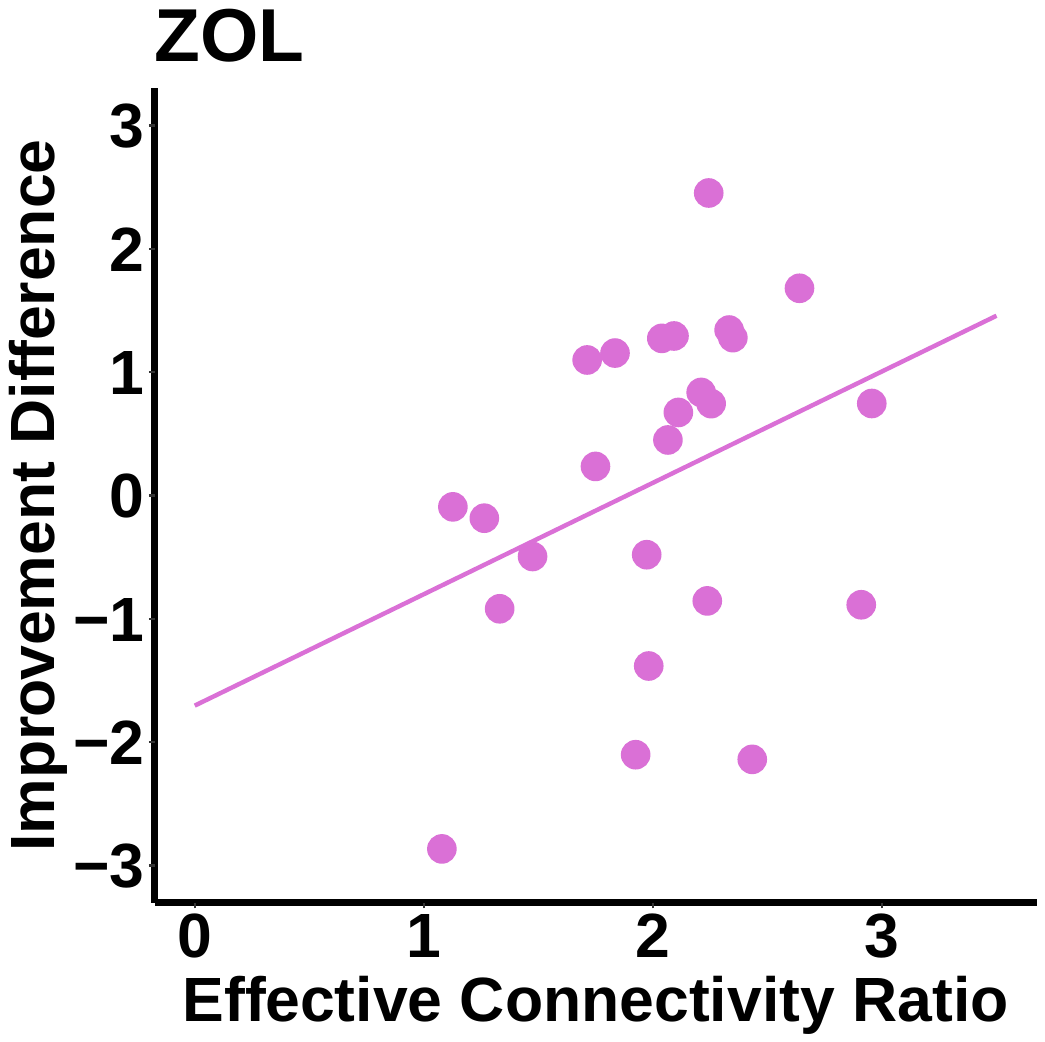

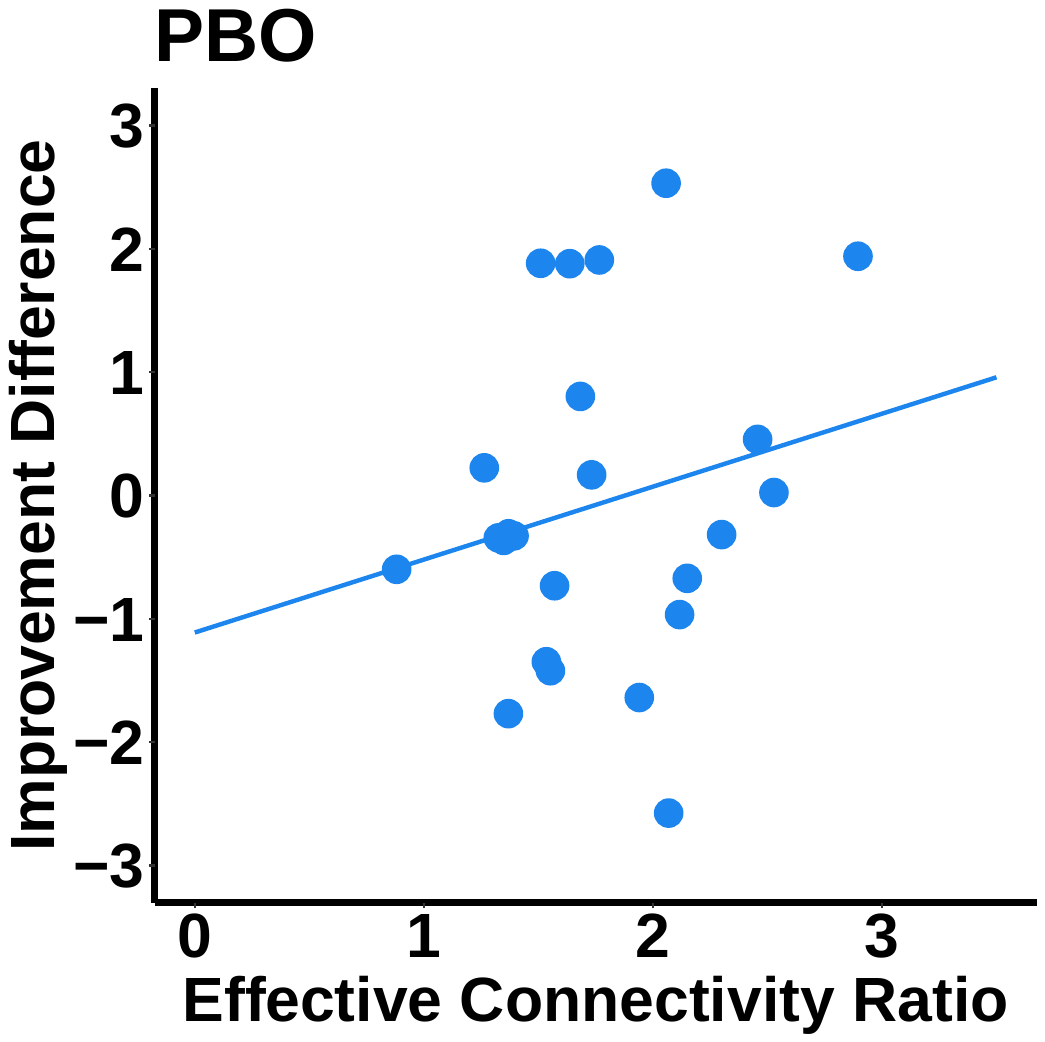

(c)

(d)

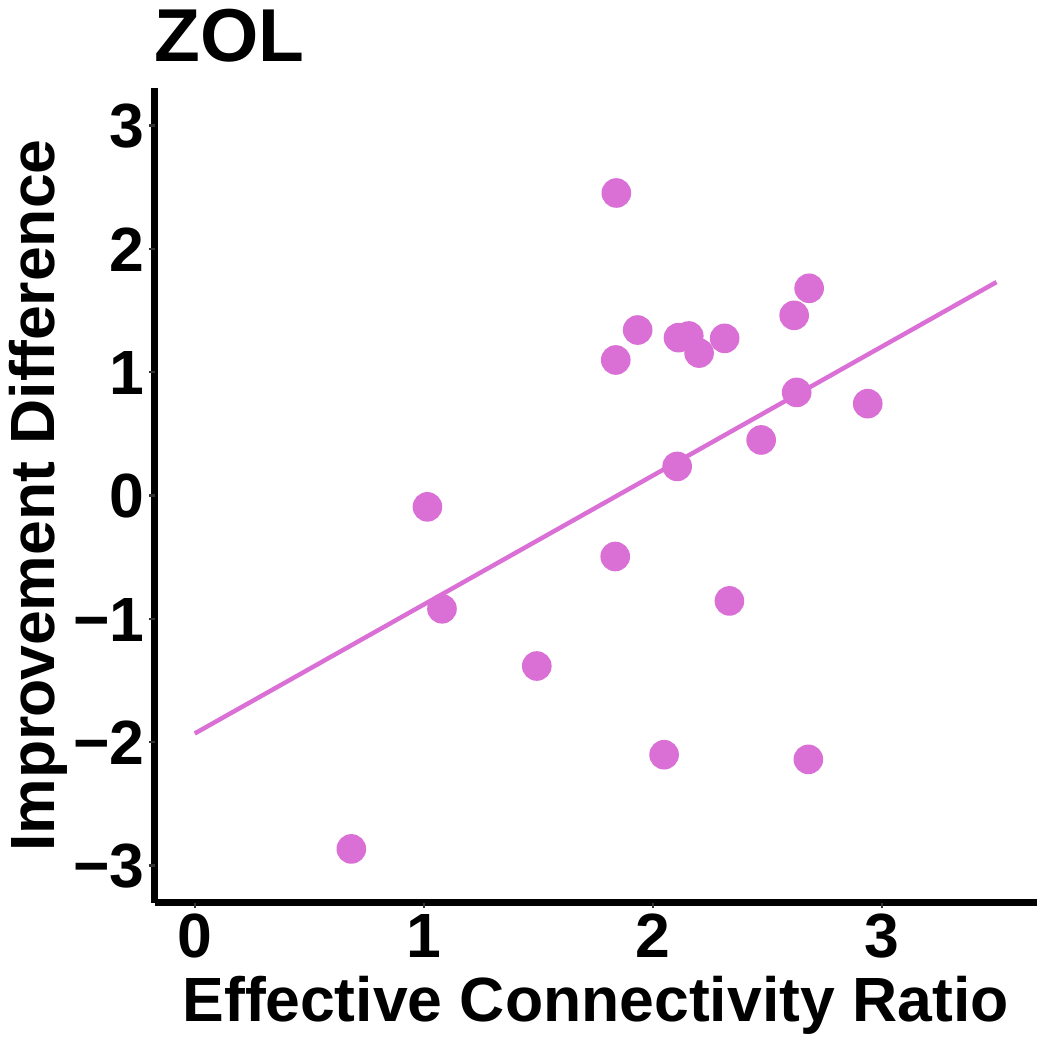

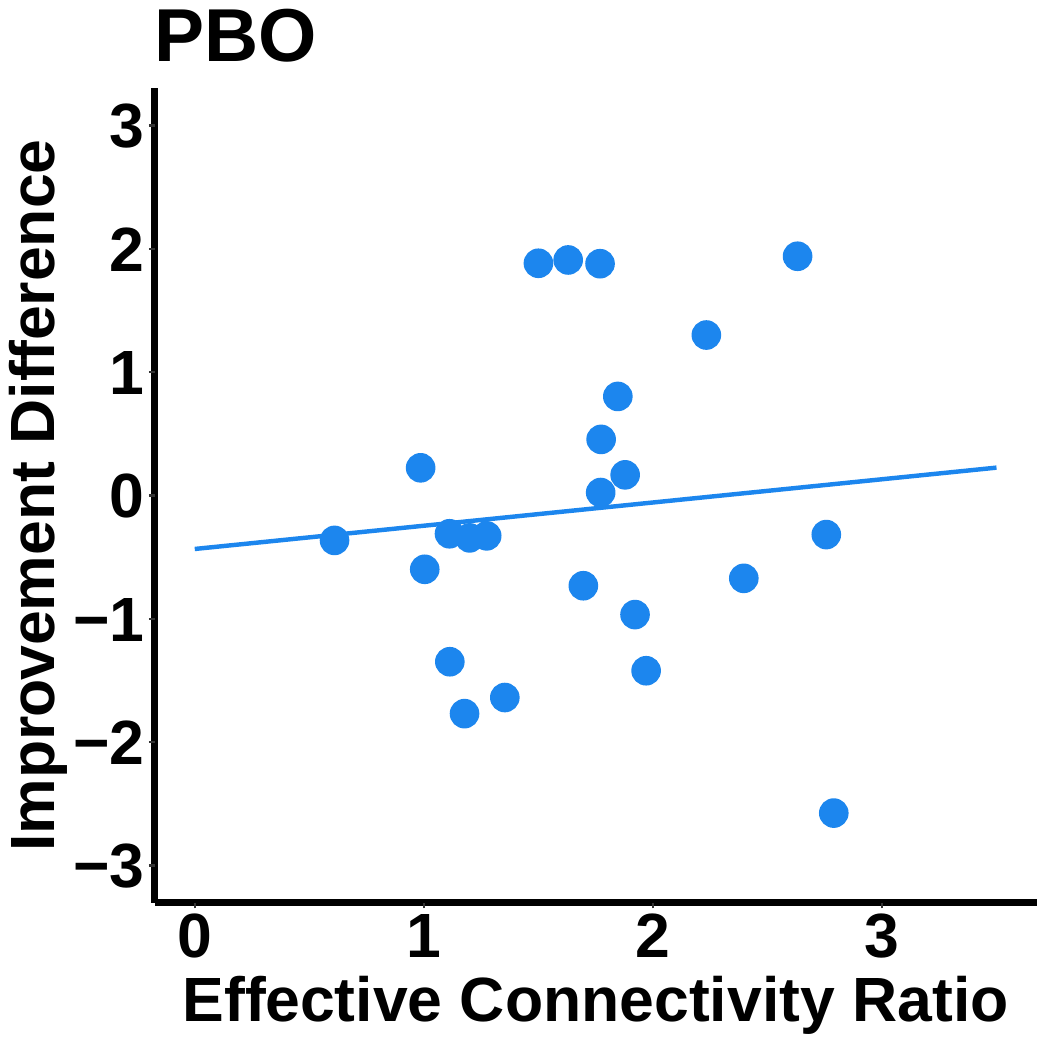

We found a trend correlation between whole-night effective connectivity ratio and memory trade-off during the zolpidem night (r = .337; p = .100; Figure A), but not the placebo night (r = .211; p = .323; Figure B). In quartile 3, we found a marginally significant correlation between effective connectivity ratio and memory trade-off during the zolpidem night (r = .427; p = .054; Figure C), but not the placebo night (r = .088; p = .682; Figure D).

Table S1. Descriptive statistics for Demographics

|  | Experiment 1 | Experiment 2 |
| --- | --- | --- |
| Age (years) | 20.38 (1.88) | 20.85 (2.97) |
| Male/female | 17/17 (50/50) | 19/19 (50/50) |
| Education (years) | 14.13 (1.66) | 14.67 (2.17) |
| ESS | 7.59 (2.81) | 6.61 (2.42) |
| BMI (kg/m^2^) | 24.29 (3.73) | 24.82 (3.45) |
| Weight (lb) | 152.38 (31.39) | 158.84 (28.17) |
| Right-handed/ left-handed | 34/0 (100/0) | 35/3 (92/8) |

*Note: Data are reported as Mean (standard deviation) for quantitative variables and N (%) for categorical variables; ESS: Epworth Sleepiness Scale; BMI: Body Mass Index.*

Table S2. Experiment 1 Sleep Architecture

| Drug | PBO | ZOL |  |
| --- | --- | --- | --- |
| TIB (min) | 531.1983 (10.0147) | 512.7203 (9.9295) | *n.s.* |
| TST (min) | 471.5086 (11.0225) | 470.7034 (10.9287) | *n.s.* |
| Stage 1 (min) | 15.0372 (1.8978) | 10.7450 (1.8899) | *n.s.* |
| Stage 2 (min) | 244.0504 (8.7200) | 234.4514 (8.7642) | *n.s.* |
| SWS (min) | 95.2869 (6.1151) | 114.6660 (6.1580) | *** |
| REM (min) | 115.2036 (5.8510) | 106.0416 (5.8623) | *n.s.* |
| WASO (min) | 37.3222 (5.7574) | 26.4467 (5.7965) | *** |
| SE (%) | 87.6834 (2.1189) | 89.7306 (2.1284) | *n.s.* |

*Note: Data are reported as Mean (standard error). TIB = Time in bed; TST = Total Sleep Time; WASO = Wake After Sleep Onset (calculated as the minutes of wake after first epoch of sleep); SE = Sleep Efficiency (calculated as 100*TST/Total Time in Bed). Asterisks indicate significant differences between conditions (n.s. p >0.05; *p < 0.05; **p < 0.01; ***p < 0.001).*

**Table S3. Experiment 2 Sleep Architecture**

| Drug | PBO | ZOL |  |
| --- | --- | --- | --- |
| TIB (min) | 582.2941 (6.2418) | 575.8939 (5.4352) | *n.s.* |
| TST (min) | 541.1471 (7.8834) | 538.4394 (7.5856) | *n.s.* |
| Stage 1 (min) | 13.4706 (1.4540) | 11.8485 (1.9770) | *n.s.* |
| Stage 2 (min) | 285.6176 (8.3986) | 284.5909 (7.6952) | *n.s.* |
| SWS (min) | 109.3676 (6.1462) | 125.4091 (7.3790) | ** |
| REM (min) | 132.4412 (5.9244) | 116.197 (5.6641) | ** |
| WASO (min) | 29.5735 (4.6177) | 24.2576 (4.2369) | *n.s.* |
| SE (%) | 92.9503 (0.9376) | 93.4436 (0.8178) | *n.s.* |

*Note: Data are reported as Mean (standard error). TIB = Time in bed; TST = Total Sleep Time; WASO = Wake After Sleep Onset (calculated as the minutes of wake after first epoch of sleep); SE = Sleep Efficiency (calculated as 100*TST/Total Time in Bed). Asterisks indicate significant differences between conditions (n.s. p >0.0; *p < 0.05; **p < 0.01; ***p < 0.001).*

**Table S4. Experiment 1 Summary of HRV Parameters Across Sleep Stages**

| Stage | Rest | |  | Stage 2 | |  | SWS | |  | REM | |  |
| --- | --- | --- | --- | --- | --- | --- | --- | --- | --- | --- | --- | --- |
| Drug/N | PBO/48 | ZOL/45 |  | PBO/54 | ZOL/54 |  | PBO/52 | ZOL/53 |  | PBO/50 | ZOL/54 |  |
| Epochs | 0.9265 (1.0750) | 0.7516 (1.1060) | n.s. | 28.3514 (1.0469) | 29.4467 (1.0195) | n.s. | 12.8394 (1.0381) | 16.7343 (1.0278) | ** | 17.1589 (1.0560) | 14.3541 (1.0195) | * |
| RR  (ms) | 943.8518 (20.0360) | 939.4200 (20.1655) | n.s. | 1019.4037 (19.9060) | 982.7458 (19.7599) | ** | 1002.7592 (19.8653) | 922.0266 (19.7976) | *** | 975.6168 (19.9479) | 970.5194 (19.7599) | n.s. |
| RMSSD (ln)  ) | 4.2359 (0.0733) | 4.2642 (0.0739) | n.s. | 4.2683 (0.0727) | 4.2194 (0.0719) | n.s. | 4.1433 (0.0725) | 3.9617 (0.0722) | *** | 4.3145 (0.0729) | 4.3611 (0.0719) | n.s. |
| HF  (ln ms^2^) | 6.6928 (0.1443) | 6.7489 (0.1455) | n.s. | 6.8488 (0.1431) | 6.7663 (0.1418) | n.s. | 6.5898 (0.1427) | 6.2717 (0.1422) | ** | 6.8766 (0.1435) | 6.9565 (0.1418) | n.s. |
| LF  (ln ms^2^) | 6.4687 (0.1066) | 6.6036 (0.1088) | n.s. | 6.6530 (0.1045) | 6.5559 (0.1024) | n.s. | 5.6299 (0.1039) | 5.6921 (0.1030) | n.s. | 6.9610 (0.1052) | 6.9894 (0.1024) | n.s. |
| TP  (ln ms^2^) | 7.7537 (0.0993) | 7.9607 (0.1009) | n.s. | 7.9888 (0.0977) | 7.9231 (0.0960) | n.s. | 7.2827 (0.0972) | 7.2369 (0.0964) | n.s. | 8.3585 (0.0982) | 8.4724 (0.0960) | n.s. |
| HF_nu_ | 0.5479 (0.0222) | 0.5279 (0.0225) | n.s. | 0.5592 (0.0220) | 0.5541 (0.0218) | n.s. | 0.7044 (0.0219) | 0.6194 (0.0218) | *** | 0.4774 (0.0221) | 0.4856 (0.0218) | n.s. |
| HF_pf_ | 0.2626 (0.0079) | 0.2607 (0.0080) | n.s. | 0.2506 (0.0078) | 0.2592 (0.0077) | n.s. | 0.2689 (0.0078) | 0.2774 (0.0077) | n.s. | 0.2276 (0.0078) | 0.2502 (0.0077) | n.s. |

*Data are reported as Mean (standard error). N: number of nights. Epochs: number of consecutive 5-min epochs per stage. Asterisks indicate significant differences between conditions (corrected by Tukey’s; n.s. p >0.05; *p < 0.05; **p < 0.01; ***p < 0.001)*

**Table S5. Experiment 2 Summary of HRV Parameters Across Sleep Stages (Q2 and Q3)**

| Stage | Stage 2 | |  | SWS | |  | REM | |  |
| --- | --- | --- | --- | --- | --- | --- | --- | --- | --- |
| Drug/ N | PBO/36 | ZOL/35 |  | PBO/35 | ZOL/33 |  | PBO/36 | ZOL/35 |  |
| Epochs | 17.92 (0.864) | 19.71 (0.876) | n.s. | 5.28 (0.963) | 4.50 (0.980) | n.s. | 11.00 (0.876) | 7.40 (0.876) | ** |
| RR (ms) | 1005 (24.8) | 960 (24.8) | *** | 980 (25.8) | 926 (26.0) | *** | 952 (24.9) | 930．(25.5) | n.s. |
| RMSSD (ln)  ) | 4.21 (.0999) | 4.04 (.0999) | ** | 4.10 (.1058) | 3.78 (.1065) | *** | 4.13 (.1003) | 4.04 (.1021) | n.s. |
| HF (ln ms^2^) | 6.81 (.187) | 6.56 (.187) | * | 6.56 (.200) | 5.96 (.201) | *** | 6.56 (.188) | 6.46 (.192) | n.s. |
| LF (ln ms^2^) | 6.43 (.126) | 6.30 (.126) | n.s. | 5.61 (.138) | 5.63 (.139) | n.s. | 6.80 (.127) | 6.66 (.131) | n.s. |
| TP (ln ms^2^) | 7.84 (.125) | 7.69 (.125) | n.s. | 7.27 (.136) | 6.97 (.137) | * | 8.20 (.125) | 8.13 (.129) | n.s. |
| HF_nu_ | .590 (.0245) | .562 (.0245) | n.s. | .692 (.0266) | .571 (.0269) | *** | .444 (.0247) | .463 (.0253) | n.s. |
| HF_pf_ | 0.242 (.00566) | 0.242 (.00567) | n.s. | 0.257 (.00647) | 0.262．(.00656) | n.s. | 0.226 (.00572) | 0.223 (.00597) | n.s. |

*Data are reported as Mean (standard error). N: number of nights. Epochs: number of consecutive 5-min epochs per stage. Asterisks indicate significant differences between conditions (corrected by Tukey’s; n.s. p >0.05; *p < 0.05; **p < 0.01; ***p < 0.001)*

**Table S6. Experiment 2 Summary of HRV Parameters Across Sleep Stages (Whole Night)**

| Stage | Rest | |  | Stage 2 | |  | SWS | |  | REM | |  |
| --- | --- | --- | --- | --- | --- | --- | --- | --- | --- | --- | --- | --- |
| Drug | PBO/36 | ZOL/35 |  | PBO/36 | ZOL/35 |  | PBO/35 | ZOL/33 |  | PBO/36 | ZOL/35 |  |
| Epochs | 1.1952  (1.3512) | 1.2298  (1.3513) | n.s. | 32.2223  (1.1923) | 34.0776  (1.2092) | n.s. | 14.9445  (1.1923) | 17.5057  (1.2267) | n.s. | 17.5834  (1.1923) | 12.9634  (1.2092) | ** |
| RR  (ms) | 921.3591  (25.57622) | 916.2846  (25.58329) | n.s. | 995.8569  (24.79799) | 965.1472  (24.87801) | * | 960.8052 (24.79799) | 912.5708 (24.9602) | ** | 952.8750 (24.7979) | 946.8405 (24.8780) | n.s. |
| RMSSD (ln)  ) | 4.116151  (0.10333661) | 4.024510  (0.10336732) | n.s. | 4.210973  (0.0999358) | 4.136072  (0.100285) | n.s. | 4.043316 (0.09993582) | 3.865868 (0.10064534) | ** | 4.164037 (0.09993582) | 4.216343 (0.10028596) | n.s. |
| HF  (ln ms^2^) | 6.548998 (0.1914384) | 6.434910 (0.1915010) | n.s. | 6.841122 (0.1844438) | 6.833192 (0.1851653) | n.s. | 6.525776 (0.1844438) | 6.211133 (0.1859056) | * | 6.692689 (0.1844438) | 6.828678 (0.1851653) | n.s. |
| LF  (ln ms^2^) | 6.358597  (0.160125) | 6.192899  (0.160206) | n.s. | 6.539154 (0.1499432) | 6.627695 (0.1510081) | n.s. | 5.596043 (0.1499432) | 5.632925 (0.1520980) | n.s. | 6.810792 (0.1499432) | 6.97889 (0.1510081) | n.s. |
| TP  (ln ms^2^) | 7.62366  (0.1964068) | 7.554683 (0.19651) | n.s. | 7.914314 (0.1817092) | 8.135149 (0.1832551) | n.s. | 7.257785  (0.1817092) | 7.126476 (0.1848365) | n.s. | 8.262321 (0.1817092) | 8.526688  (0.1832551) | n.s. |
| HF_nu_ | 0.5366224 (0.02570365) | 0.5547556 (0.02571716) | n.s. | 0.5696665 (0.02399774) | 0.5661786 (0.02417641) | n.s. | 0.6875529 (0.02399774) | 0.6143552 (0.02435925) | ** | 0.4654334 (0.02399774) | 0.4694941 (0.02417641) | n.s. |
| HF_pf_ | 0.2655057 (0.008260159) | 0.2477035 (0.008264036) | n.s. | 0.2396786 (0.007401850) | 0.2397315 (0.007493175) | n.s. | 0.2596796 (0.007401850) | 0.2625183 (0.007586984) | n.s. | 0.2219946 (0.007401850) | 0.2257583 (0.007493175) | n.s. |

*Data are reported as Mean (standard error). Epochs: number of consecutive 5-min epochs per stage. Asterisks indicate significant differences between conditions (corrected by Tukey’s; n.s. p >0.05; *p < 0.05; **p < 0.01; ***p < 0.001)*

**Table S7. Experiment 1 Normalized EEG Power during NREM Modulated by Drug Conditions (Whole Night)**

| Stage | | Stage 2 | |  | SWS | |  |
| --- | --- | --- | --- | --- | --- | --- | --- |
| Drug | | PBO | ZOL |  | PBO | ZOL |  |
| SWA | Frontal | .2938 (.0161) | .3063 (.0169) | n.s. | .4436 (.0209) | .4637 (.0200) | n.s. |
| 0.5-2Hz | Central | .2668 (.0142) | .2821 (.0162) | n.s. | .4104 (.0201) | .4409 (.0203) | n.s. |
|  | Parietal | .2539 (.0139) | .2647 (.0159) | n.s. | .3968 (.0206) | .4286 (.0214) | n.s. |
|  | Occipital | .2212 (.0138) | .2387 (.0158) | n.s. | .3437 (.0212) | .3887 (.0225) | n.s. |
| Delta | Frontal | .1966 (.0135) | .1952 (.0137) | n.s. | .2550 (.0150) | .2329 (.0131) | n.s. |
| 1-4Hz | Central | .1895 (.0128) | .1900 (.0135) | n.s. | .2294 (.0145) | .2159 (.0131) | n.s. |
|  | Parietal | .1832 (.0123) | .1807 (.0127) | n.s. | .2163 (.0142) | .2014 (.0127) | n.s. |
|  | Occipital | .1651 (.0120) | .1723 (.0124) | n.s. | .1806 (.0134) | .1812 (.0119) | n.s. |
| Sigma | Frontal | .0123 (.0017) | .0149 (.0013) | n.s. | .0059 (.0015) | .0059 (.0005) | n.s. |
| 12-16Hz | Central | .0166 (.0018) | .0212 (.0018) | * | .0073 (.0016) | .0078 (.0006) | n.s. |
|  | Parietal | .0223 (.0022) | .0274 (.0022) | * | .0088 (.0016) | .0095 (.0008) | n.s. |
|  | Occipital | .0180 (.0019) | .0223 (.0019) | * | .0076 (.0016) | .0079 (.0007) | n.s. |
| Theta | Frontal | .0439 (.0045) | .0370 (.0029) | n.s. | .0327 (.0037) | .0241 (.0014) | n.s. |
| 4-8Hz | Central | .0518 (.0048) | .0432 (.0032) | n.s. | .0344 (.0038) | .0256 (.0015) | n.s. |
|  | Parietal | .0580 (.0049) | .0477 (.0035) | n.s. | .0368 (.0039) | .0275 (.0018) | n.s. |
|  | Occipital | .0664 (.0058) | .0587 (.0044) | n.s. | .0432 (.0042) | .0360 (.0026) | n.s. |

*Data are reported as Mean (standard error). Powers were averaged bilaterally from channel F3, F4, C3, C4, P3, P4, O1, O2. (Asterisks indicate significant differences between drug conditions by linear-mixed models; n.s. p >0.05; *p < 0.05; **p < 0.01; ***p < 0.001).*

**Table S8. Experiment 2 Normalized EEG Power during NREM Modulated by Drug Conditions (Q2 & Q3)**

| Stage | | Stage 2 | |  | SWS | |  |
| --- | --- | --- | --- | --- | --- | --- | --- |
| Drug | | PBO | ZOL |  | PBO | ZOL |  |
| SWA | Frontal | .4683 (.0107) | .4723 (.0133) | n.s. | .6254 (.0148) | .6188 (.0181) | n.s. |
| 0.5-2Hz | Central | .4216 (.0109) | .4271 (.0131) | n.s. | .5898 (.0134) | .5849 (.0178) | n.s. |
|  | Parietal | .4039 (.0098) | .4030 (.0129) | n.s. | .5888 (.0138) | .5746 (.0187) | n.s. |
|  | Occipital | .3602 (.0113) | .3660 (.0142) | n.s. | .5199 (.0167) | .5087 (.0223) | n.s. |
| Delta | Frontal | .3117 (.0103) | .3041 (.0113) | n.s. | .3501 (.0114) | .3057 (.0125) | ** |
| 1-4Hz | Central | .2970 (.0099) | .2958 (.0106) | n.s. | .3210 (.0108) | .2829 (.0126) | ** |
|  | Parietal | .2898 (.0091) | .2870 (.0106) | n.s. | .3158 (.0092) | .2740 (.0126) | ** |
|  | Occipital | .2735 (.0114) | .2772 (.0121) | n.s. | .2863 (.0104) | .2531 (.0132) | * |
| Sigma | Frontal | .0238 (.0019) | .0286 (.0026) | ** | .0096 (.0010) | .0121 (.0015) | * |
| 12-16Hz | Central | .0314 (.0023) | .0394 (.0031) | *** | .0125 (.0011) | .0161 (.0018) | * |
|  | Parietal | .0395 (.0030) | .0497 (.0039) | *** | .0150 (.0014) | .0197 (.0022) | * |
|  | Occipital | .0330 (.0022) | .0401 (.0031) | ** | .0130 (.0012) | .0164 (.0018) | n.s. |
| Theta | Frontal | .0693 (.0037) | .0605 (.0037) | *** | .0436 (.0026) | .0353 (.0025) | ** |
| 4-8Hz | Central | .0816 (.0041) | .0724 (.0040) | ** | .0492 (.0031) | .0402 (.0030) | ** |
|  | Parietal | .0897 (.0043) | .0785 (.0042) | ** | .0548 (.0035) | .0443 (.0034) | ** |
|  | Occipital | .1046 (.0056) | .0944 (.0056) | n.s. | .0703 (.0049) | .0585 (.0051) | * |

*Data are reported as Mean (standard error). Powers were averaged bilaterally from channel F3, F4, C3, C4, P3, P4, O1, O2. (Asterisks indicate significant differences between drug conditions by paired t-tests; n.s. p >0.05; *p < 0.05; **p < 0.01; ***p < 0.001).*

**Table S9. Correlations between HRV Parameters and Behavioral Improvements**

|  | WPA | | | | | | OS | | | | | |
| --- | --- | --- | --- | --- | --- | --- | --- | --- | --- | --- | --- | --- |
|  | HFln | | RMSSDln | | HFnu | | HFln | | RMSSDln | | HFnu | |
| Drug | PBO | ZOL | PBO | ZOL | PBO | ZOL | PBO | ZOL | PBO | ZOL | PBO | ZOL |
| Stage 2 | r = -.035  p = .851 | r = -.460  p = .008 | r = -.094  p = .615 | r = -.508  p = .003 | r = -.036  p = .846 | r = -.406  p = .021 | r = .051  p = .773 | r = .015  p = .933 | r = .086  p = .629 | r = -.065  p = .709 | r = -.019  p = .915 | r = .117  p = .501 |
| SWS  ) | r = -.096  p = .654 | r = -.460  p = .018 | r = -.168  p = .431 | r = -.513  p = .007 | r = -.014  p = .948 | r = -.350  p = .080 | r = .422  p = .032 | r = .131  p = .507 | r = .397  p = .044 | r = .068  p = .731 | r = .141  p = .491 | r = .220  p = .260 |

*Data are reported as Pearson’s correlation coefficients (r) and p-values (p).*

**Table S10. SOs Counts and Sigma/SOs by Drug Conditions**

| Drug | | PBO | ZOL |  |
| --- | --- | --- | --- | --- |
| SO | F3 | 169.242 (28.061) | 160.909 (44.758) | n.s. |
| Counts | F4 | 185.242 (34.602) | 174.484 (49.172) | n.s. |
|  | C3 | 122.636 (20.164) | 105.485 (25.098) | n.s. |
|  | C4 | 102.970 (22.061) | 94.333 (22.011) | n.s. |
|  | P3 | 104.212 (18.445) | 78.757 (16.317) | * |
|  | P4 | 94.394 (18.007) | 71.303 (16.093) | * |
| SO up-state | F3 | 0.01066241 (0.000572956) | 0.01052726 (0.0007072569) | n.s. |
| Sigma | F4 | 0.01179984 (0.0009264091) | 0.01172965 (0.0007314609) | n.s. |
| Normalized | C3 | 0.01175520 (0.0006375368) | 0.01146652 (0.0007709227) | n.s. |
| Power | C4 | 0.01066014 (0.0007590402) | 0.01076715 (0.0006134255) | n.s. |
|  | P3 | 0.01034928 (0.0006057417) | 0.01170495 (0.0008417385) | n.s. |
|  | P4 | 0.010588097 (0.0006876312) | 0.009728913 (0.0006714047) | n.s. |

*Data are reported as Mean (standard error). (Asterisks indicate significant differences between drug conditions by paired t-tests; n.s. p >0.05; *p < 0.05).*

**Table S11. Correlations between Sigma/SOs couplings and Behavioral Improvements**

| PBO | F3 | F4 | C3 | C4 | P3 | P4 |
| --- | --- | --- | --- | --- | --- | --- |
| WPA | r = .33  p = .082 | r = .28  p = .150 | r = .24  p = .200 | r = .34  p = .073 | r = .40  p = .034 | r = .21  p = .27 |
| OS | r = -.51  p = .003 | r = -.33  p = .075 | r = -.31  p = .085 | r = -.17  p = .370 | r = -.38  p = .033 | r = -.16  p = .400 |
| WPA - OS  ) | r = .59  p < .001 | r = .45  p = .016 | r = .39  p = .036 | r = .38  p = .041 | r = .560  p = .002 | r = .29  p = .130 |

| ZOL | F3 | F4 | C3 | C4 | P3 | P4 |
| --- | --- | --- | --- | --- | --- | --- |
| WPA | r = .16  p = .378 | r = -.11  p = .543 | r = .16  p = .373 | r = -.18  p = .351 | r = .12  p = .530 | r = -.09  p = .632 |
| OS | r = .20  p = .291 | r = -.04  p = .817 | r = .11  p = .559 | r = -.18  p = .351 | r = .24  p = .217 | r = -.14  p = .472 |
| WPA - OS  ) | r = -.10  p = .598 | r = -.60  p = .758 | r = -.02  p = .915 | r = -.01  p = .950 | r = -.12  p = .551 | r = .02  p = .911 |

*Data are reported as Pearson’s correlation coefficients (r) and p-values (p).*

**Table S12. Descriptive statistics for Behavioral Tasks**

|  |  | Test 1 | Test 2 | Test 3 |
| --- | --- | --- | --- | --- |
| WPA | PBO | 0.816 (0.238) | 0.556 (0.290) | 0.442 (0.273) |
|  | ZOL | 0.840 (0.200) | 0.583 (0.275) | 0.513 (0.253) |
| OS | PBO | 0.618 (0.171) | 0.661 (0.152) | 0.696 (0.148) |
|  | ZOL | 0.633 (0.149) | 0.652 (0.155) | 0.663 (0.161) |

*Data are reported as Mean (standard deviation).*

**Table S13. Effective Connectivity Window Numbers (Each Window Consists of 500 samples of 5-second sliding windows)**

|  | All Stages | NREM |
| --- | --- | --- |
| Study 1 PBO | 23.0385 (3.0798) | 9.3654 (3.6283) |
| Study 1 ZOL | 22.5769 (3.5311) | 9.2115 (3.4981) |
| Study 2 PBO | 25.7273 (2.0657) | 8.9697 (2.6278) |
| Study 2 ZOL | 25.1515 (1.9862) | 10.2727 (2.6725) |

*Data are reported as Mean (standard deviation). Note: In the methods, we mentioned that the MVAR and GPDC were computed for each window with length of 500 samples of 5-second sliding windows, which is 500 * 5 s = 2500 seconds. Here, we excluded REM and WASO by removing windows that consists of any REM or WASO. For example, if the 300th sample include REM, we exclude the first 500 samples and the second 500 samples. This step dropped our window numbers from ~24 to ~9.*
